# Discovery of re-purposed drugs that slow SARS-CoV-2 replication in human cells

**DOI:** 10.1101/2021.01.31.428851

**Authors:** Adam Pickard, Ben C. Calverley, Joan Chang, Richa Garva, Yinhui Lu, Karl E. Kadler

## Abstract

COVID-19 vaccines based on the Spike protein of SARS-CoV-2 have been developed that appear to be largely successful in stopping infection. However, vaccine escape variants might arise leading to a re-emergence of COVID. In anticipation of such a scenario, the identification of repurposed drugs that stop SARS-CoV-2 replication could have enormous utility in stemming the disease. Here, using a nano-luciferase tagged version of the virus (SARS-CoV-2- DOrf7a-NLuc) to quantitate viral load, we evaluated a range of human cell types for their ability to be infected and support replication of the virus, and performed a screen of 1971 FDA-approved drugs. Hepatocytes, kidney glomerulus, and proximal tubule cells were particularly effective in supporting SARS-CoV-2 replication, which is in- line with reported proteinuria and liver damage in patients with COVID-19. We identified 35 drugs that reduced viral replication in Vero and human hepatocytes when treated prior to SARS-CoV-2 infection and found amodiaquine, atovaquone, bedaquiline, ebastine, LY2835219, manidipine, panobinostat, and vitamin D3 to be effective in slowing SARS-CoV-2 replication in human cells when used to treat infected cells. In conclusion, our study has identified strong candidates for drug repurposing, which could prove powerful additions to the treatment of COVID.

## INTRODUCTION

The COVID-19 pandemic caused by the severe acute respiratory syndrome coronavirus 2 (SARS-CoV-2) virus is having a widespread impact on global health with substantial loss of life. SARS-CoV-2 infection in patients with COVID-19 can result in pulmonary distress, inflammation, and broad tissue tropism. Some infected individuals are asymptomatic whilst others mount an exaggerated immune response, or ‘cytokine storm’(Chatenoud *et al*, 1991; Fajgenbaum & June, 2020), which correlates with multiple organ failure and poor outcome (Ruan *et al*, 2020). Vaccines have been developed to help protect people from COVID-19 but the emergence of mutational variants, some with increased transmissibility, calls for additional research into alternative treatment and prevention strategies.

An early step in the infection process is interaction of the Spike protein on the surface of the virus with angiotensin-converting enzyme 2 (ACE2) on the surface of the host cell (Li *et al*, 2003). Using this information, first-generation vaccines have been generated against the Spike protein. SARS-CoV-2 entry can be potentiated by additional host factors including neuropilin-1 (Cantuti-Castelvetri *et al*, 2020). Once inside, the virus uses components of the host cell to replicate and secrete viral particles (Maier *et al*, 2016) and disrupt RNA handling and protein translation to suppress host defenses (Banerjee *et al*, 2020). The different stages of the disease, from the initial infection of host cells through to virus replication and the response (normal or extreme) of the immune system, offer opportunities to identify drugs, treatments and therapies to help stop disease progression. Furthermore, large proportions of the world’s population remain at risk of contracting COVID-19 as they wait to be vaccinated, which has prompted governments to encourage people to continue to self-isolate, remain at home, and maintain social distancing. The identification of safe and easily distributed medications that can target the different stages of virus infection and replication, could reduce the spread of SARS-CoV-2 and reduce the cases of COVID-19.

The Food and Drug Administration (FDA) and the European Medicines Agency (EMA) work with pharmaceutical companies to develop safe and effective drugs for the benefit of public health. Using a library of 1971 FDA-approved compounds, here we used a traceable clone of SARS-Cov-2 to identify well-characterized drugs that could potentially slow the infection and replication of the SARS-CoV-2 virus in human cells.

We had previously developed an approach to quantitate collagen synthesis by using CRISPR-Cas9 to insert the gene encoding NanoLuciferase (NLuc) into the Col1a2 gene in fibroblasts (Calverley *et al*, 2020). Here, we adapted the approach to tag the SARS-CoV-2 virus with NLuc, and then used the recombinant NLuc-tagged virus to monitor virus replication and to identify drugs that slow the replication process. SARS-CoV-2 was originally recovered by culturing human specimens in the African green monkey kidney cell line, Vero (Zhou *et al*, 2020), and the virus readily replicates in this cell type. We have performed initial screening in the Vero cell line, and in order to draw reliable conclusions from our studies we performed a replicate screen in a human hepatocyte cell line, HUH7. We evaluated a panel of human cell types for their ability to be infected by SARS-CoV-2 and to support virus replication, and optimized culture conditions for each cell type to provide a robust system for screening in human cells. We found that hepatocytes and kidney epithelial cells were proficient in supporting SARS-CoV-2 replication whereas fibroblasts and lung epithelial cells supported only minimal replication.

Through applying stringent criteria in our hit selection and confirming efficacy of these compounds in suppressing replication in cells already infected with SARS-CoV-2, we identified a shortlist of nine drugs that were effective in inhibiting SARS-CoV-2 replication. The drugs identified included anti-cancer and anti-viral drugs, but also less toxic drugs such as atovoquone, ebastine and vitamin D3. The multiple steps involved in virus infection and proliferation, means that each of these drugs may have different molecular targets, and for some, additional targets are still being elucidated. As these drugs are FDA-approved and safe, and dosimetry has been established for use in people, clinical trials could be initiated for these drugs.

## RESULTS

### Generation of a traceable SARS-CoV-2 virus

High throughput screens have been developed to identify drugs suitable for re-purposing for treatment of COVID19 (e.g., (Chen *et al*, 2020; Riva *et al*, 2020)). These screens have been performed in non-human cell lines, such as Vero, and rely on secondary factors such as cell viability to identify candidates, or markers that indicate virus infection. To monitor the replication of SARS-CoV-2 we have generated a modified traceable virus where Orf7a is replaced with the sequence encoding NanoLuciferase (SARS-Cov-2-ΔOrf7a-NLuc, **Fig. 1A**). NanoLuciferase (NLuc) is an enzyme that produces light when supplied with its substrate (coelenterazine) and is readily detectable even at low quantities (Calverley *et al*., 2020). Orf7a has previously been removed in SARS-CoV and SARS-CoV-2 and yielded infectious and replicative virus particles (Thi Nhu Thao *et al*, 2020; Xie *et al*, 2020a; Xie *et al*, 2020b). We have used a reverse-genetics approach to generate virus particles (**Supplemental Fig. 1**), and plaque forming assays demonstrated that the replication of SARS-CoV-2-ΔOrf7a-NLuc viruses were equivalent to the wild-type SARS-CoV-2 (Wuhan-Hu-1, NC_045512.2) (**Fig. 1B**). Viral RNAs (**Supplemental Fig. 1**), and electron microscopy (**Fig. 1C-D, Supplementary Fig. 2**) confirmed virus production. NLuc activity was readily detected in infected cultures upon addition of the substrate coelentrazine (**Fig. 1E**).

**Figure 1:**
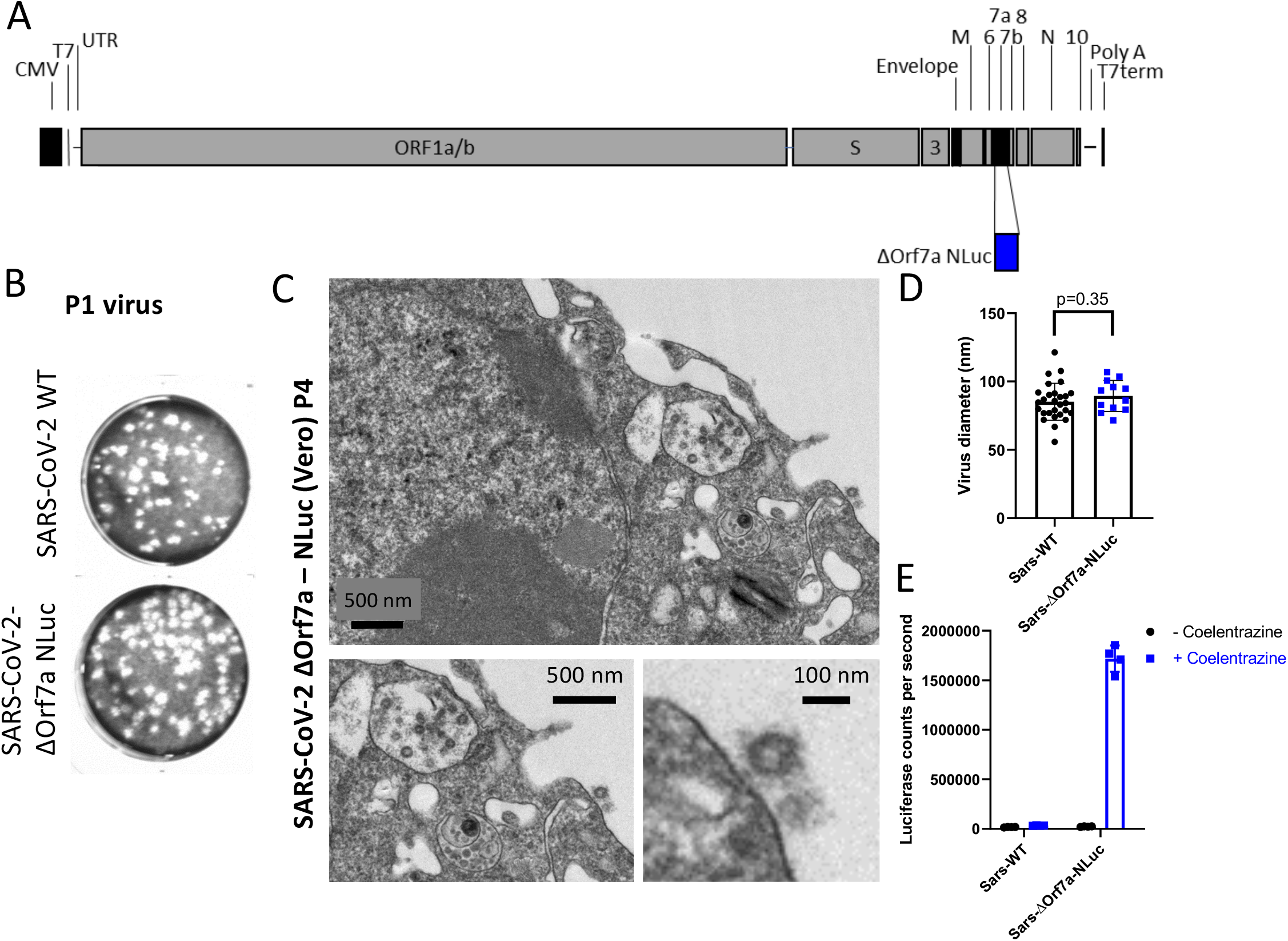
Nanoluciferase (NLuc) modified SARS-CoV-2 virus as a reporter for virus replication. A) Diagram of the SARS-CoV-2 genome highlighting the insert site for the reporter NLuc in place of Orf7a, (SARS-CoV-2- ΔOrf7a-Nluc). B) Virus particles recovered following transfection of wild-type (WT) or NLuc modified SARS-CoV-2 encoding RNA into 293T cells were used to infect Vero cells. The recovered virus (P1 virus) was then titred in Vero cells to assess virus replication and plaque forming potential. C) Electron microscopy of the SARS-Cov-2 virus 72 h.p.i. of Vero cells (Passage 4, P4), both intra- and extra-cellular virus particles were identified. D) Measurement of diameter for the wild-type (WT) and NLuc modified SARS-CoV-2 particles.

### NLuc activity as a marker of virus replication

To monitor replication of the SARS-CoV-2-Δ Orf7a-NLuc virus, Vero cells were exposed to increasing numbers of replicative virus particles (plaque forming units, PFU) and luminescence was used as a measure of virus replication. Luminescence measurements were compared to SARS-CoV-2-Δ Orf7a-NLuc virus in the absence of cells as background (**Fig. 2A**), 24, 48 and 72 hours post infection (h.p.i.). A lag in virus replication of at least 48 hrs was observed but replication was readily observed at 72 hrs, when as little as 2 PFU per well (MOI 0.0004) was added. Using these optimization experiments 72 h.p.i. was chosen as the time point at which to assess virus replication in subsequent assays, the average Z’ factor (Zhang *et al*, 1999) for our replication assay in 96 well format was 0.8 indicating suitability for library screening. Amplification of NLuc signal upon virus infection was consistent for all passages of the traceable virus (**Supplementary Fig. 3**) and with varying cell number (**Supplementary Fig. 4**).

**Figure 2:**
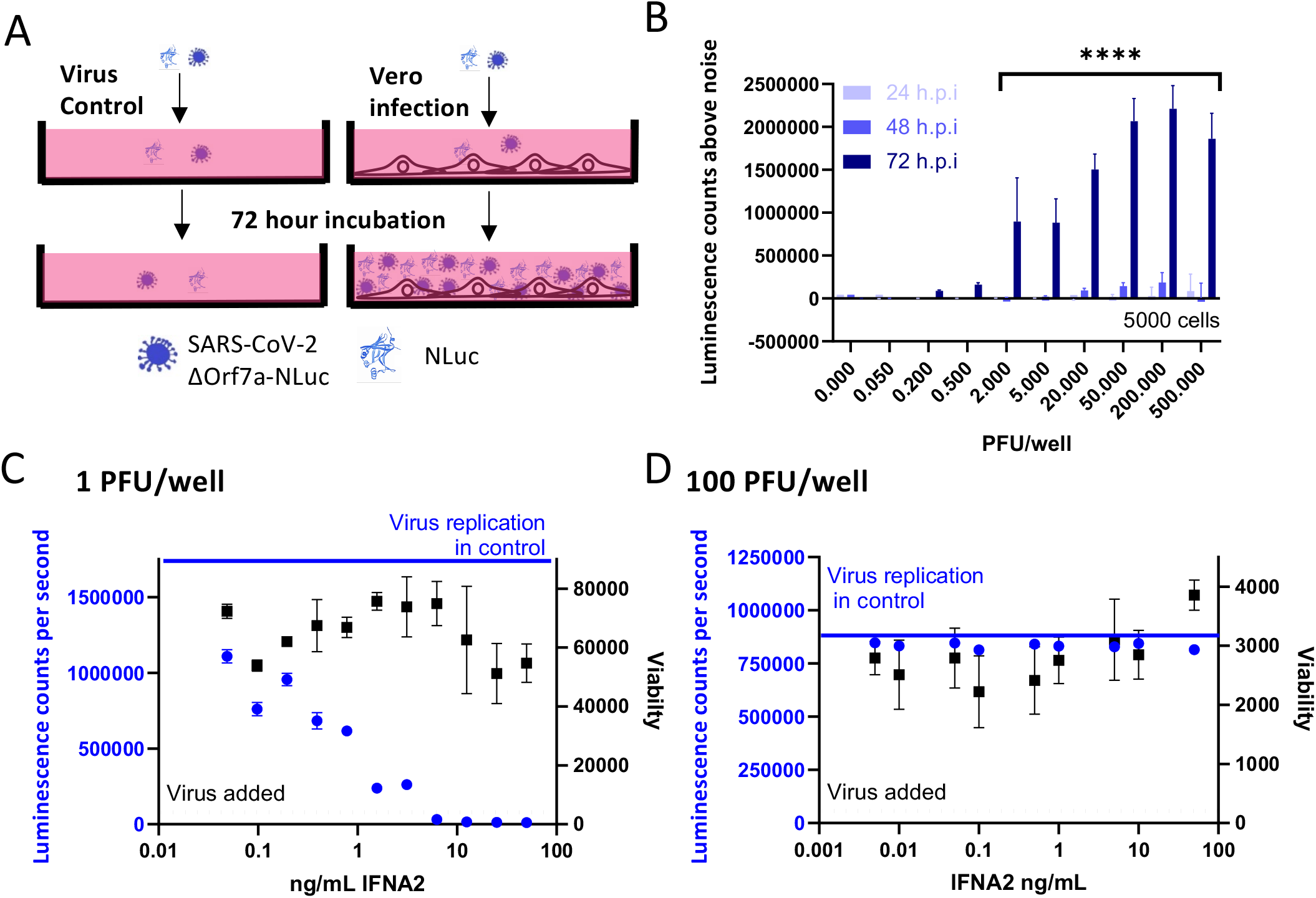
Timings of SARS-CoV-2-ΔOrf7a-NLuc virus replication. A) Schematic showing how SARS-CoV-2 ΔOrf7a-NLuc was used to assess virus replication. During virus replication as virus particles are released from the cell, or as a result of cell death, NLuc activity is detected in the conditioned medium together with virus particles. In subsequent assays the NLuc activity is recorded in the absence of cells, and NLuc signals above background signal are used to indicate the level of virus replication. B) The replication of SARS-CoV-2-ΔOrf7a-Nluc was monitored in a 96 well assay. Using 5000 Vero cells per well, increasing numbers of virus particles were added and monitored 24, 48 and 72 hours post infection (h.p.i). Substantial and significant increases in NLuc activity were observed when only two plaque forming units (PFU) were added. Higher NLuc activity was associated with higher viral input. Even with higher viral inputs the substantial increases in NLuc activity occurred only after 72 h.p.i. C) Five thousand Vero cells were pre-treated with increasing doses of IFNα2 for 24 hours prior to infection with 1PFU SARS-CoV-2-ΔOrf7a-Nluc. NLuc activity and viability were assessed 72 h.p.i. demonstrating effective inhibition of virus replication. The blue line indicates the luminescence counts per second for DMSO treated controls. N=3 replicate samples D) As in C, 5000 Vero cells were treated with increasing doses of IFNα2 for 24 hours prior to infection with 100PFU SARS- CoV-2-ΔOrf7a-Nluc. With this higher titre, virus replication was not inhibited. The blue line indicates the luminescence counts per second for DMSO treated controls. N=3 replicate samples.

Earlier studies have demonstrated interferon alpha 2 (IFNA2) is able to reduce SARS-CoV-2 replication (Xie *et al*., 2020a). We confirmed this observation using our assay (**Fig. 2C**) but noted that with higher virus load (100 PFU), IFNA2 had little effect on virus replication (**Fig. 2D**). These results provided confidence that the traceable virus could be used to quantify the effects of drugs on virus replication.

### SARS-CoV-2 replication screen validation

High throughput screens have been developed to identify drugs suitable for re-purposing for treatment of COVID19 (e.g., (Jeon *et al*, 2020; Riva *et al*., 2020; Touret *et al*, 2020)). These screens have been performed in non-human cell lines, such as Vero, and rely on secondary factors such as cell viability to identify candidates, or markers that indicate virus infection. A first step in validating our viral replication screen was to measure virus replication after treatment with interferon alpha 2 (IFNA2) which has previously demonstrated to reduce virus replication (Xie *et al*., 2020a). When Vero cells were infected with a single PFU, pre-treatment of IFNA2 was able to reduce replication of SARS-CoV-2-ΔOrf7a-NLuc in a dose dependent manner. However, with higher virus load (100 PFU), IFNA2 had little effect on virus replication (**Fig. 2C and D**). These results provided confidence that the NLuc-tagged virus system could be used to quantify the effects of drugs on virus replication, especially if reduction of bioluminescence was detected with high PFU.

### Drug repurposing to target SARS-CoV-2 replication

A 1971 FDA-approved compound library was used to pretreat Vero cells before infection with either 1PFU (MOI 0.0002) or 100 PFU (MOI 0.02) SARS-CoV-2-Δ Orf7a-NLuc virus as outlined in **Fig. 3A**. Infected Vero cells that were untreated and background (virus only wells) were used as controls. The Z’ factor was assessed for each plate of the screen averaging 0.867± 0.140 SD and 0.915± 0.058 SD, for the 1PFU and 100PFU screen, respectively demonstrating a robust assay. More hits were identified in the 1PFU screen and therefore we focused our hit selection on compounds that reduced virus replication by at least 85% in the 100PFU screen (**Fig. 3B** and **Supplementary Table 3**). We identified 69 compounds that reduced replication (FDR<0.1 compared to untreated and DMSO controls, 55 showed no viral replication). These hits were also identified in the duplicate screen performed with 1PFU per well (**Fig. 3C**). Five of these compounds were false positive hits due to direct inhibition of NLuc activity (**Fig. 3C, Supplemental Table 4**).

**Figure 3:**
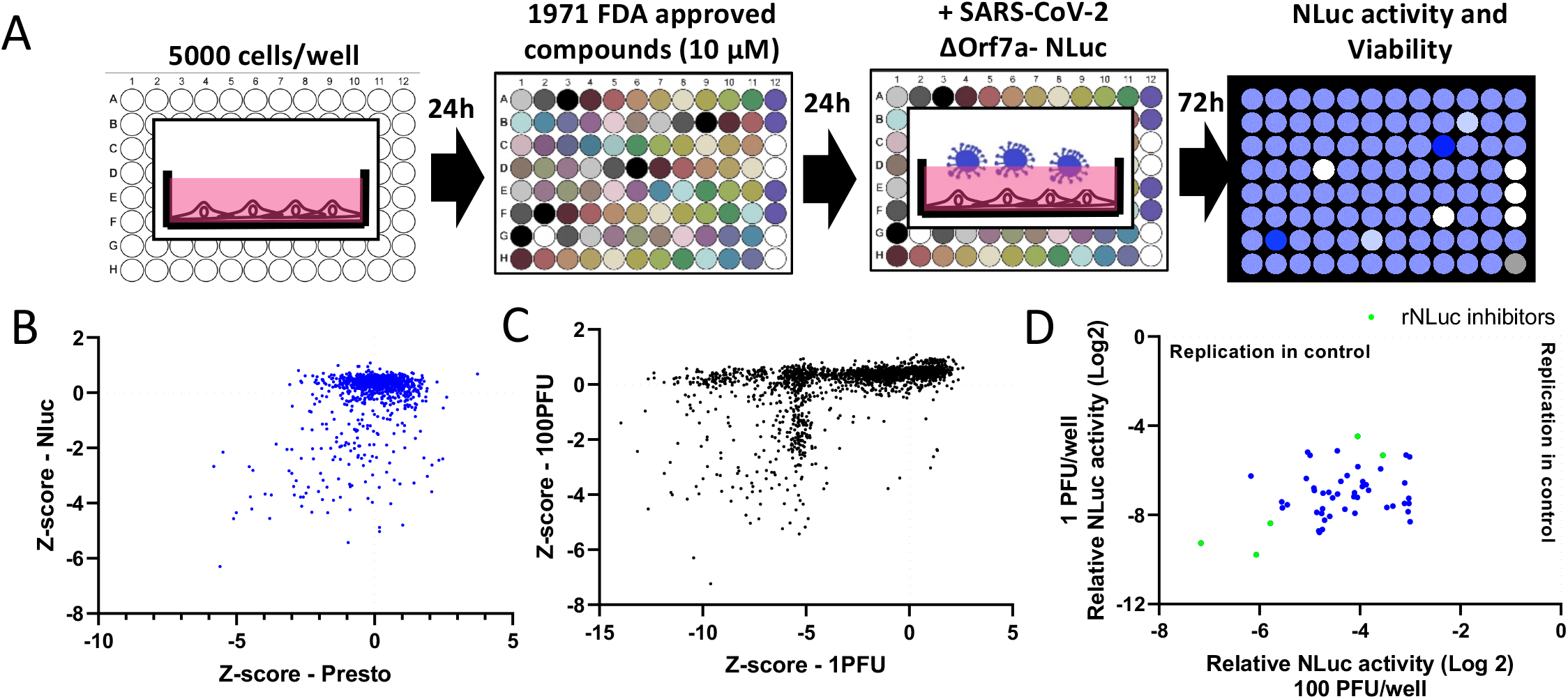
Screen of 1971 FDA-approved compounds to identify therapeutics that inhibit SARS-CoV-2-ΔOrf7a-NLuc virus replication. A) Schematic of the screening procedure used to assess whether FDA-approved compounds alter SARS-CoV-2 ΔOrf7a- Nluc virus replication. B) Scatterplot showing the effects of 1971 compounds on Sars-CoV-2 ΔOrf7a-NLuc replication and cell viability. C) Scatterplot showing the effects of 1971 compounds on Sars-CoV-2 ΔOrf7a-NLuc replication in replicate screens using 100 PFU or 1 PFU per well. The Z-score for luciferase activity is plotted. D) Scatterplot of 50 compounds that reduced NLuc activity in C. The luciferase activities relative to untreated controls (indicated by the blue dashed line) are shown. Highlighted in green are compounds that inhibit the NLuc enzyme directly.

### Drug repurposing screen for SARS-CoV-2 replication in human cells

Having established an effective screening strategy in Vero cells, we sought to establish a similar approach in a human cell line capable of supporting SARS-CoV-2 replication. SARS-CoV-2 has been reported to infect multiple cell lines, and replication of the viral genome has been identified (Chu *et al*, 2020). We tested the replicative capacity of SARS-CoV-2-Δ Orf7a in the lung epithelial cell lines Calu3, A549, and 16HBEo and in lung fibroblasts. Using real-time PCR and western blotting for the SARS-CoV-2 nucleocapsid protein, we observed virus replication in these cell lines, and we confirmed that SARS-CoV-2 WT and SARS-CoV-2-ΔOrf7a-NLuc viruses infected these lung cell lines (**Supplemental Fig. 5**). However the Z’ factor suggested that they were not suitable for high throughput screening (**Fig. 4A**). We assessed additional cell lines reported to be infected by SARS-CoV-2 including Caco-2 (Chu *et al*., 2020; Touret *et al*., 2020) and HUH7 (Zhou *et al*., 2020), as well as additional kidney cell lines. These were chosen as we were able to recover virus particles from 293T cells and kidney is reported to express high levels of ACE2 (Wysocki *et al*, 2020). Replication in monocytic cells and fibroblastic cell lines was also monitored due to the systemic and fibrotic nature of COVID-19. We identified virus replication in HUH7 cells, microvascular cells of the kidney glomerulus, and proximal tubule cells of the kidney and THP1 monocytic cells (**Fig. 4A**).

**Figure 4:**
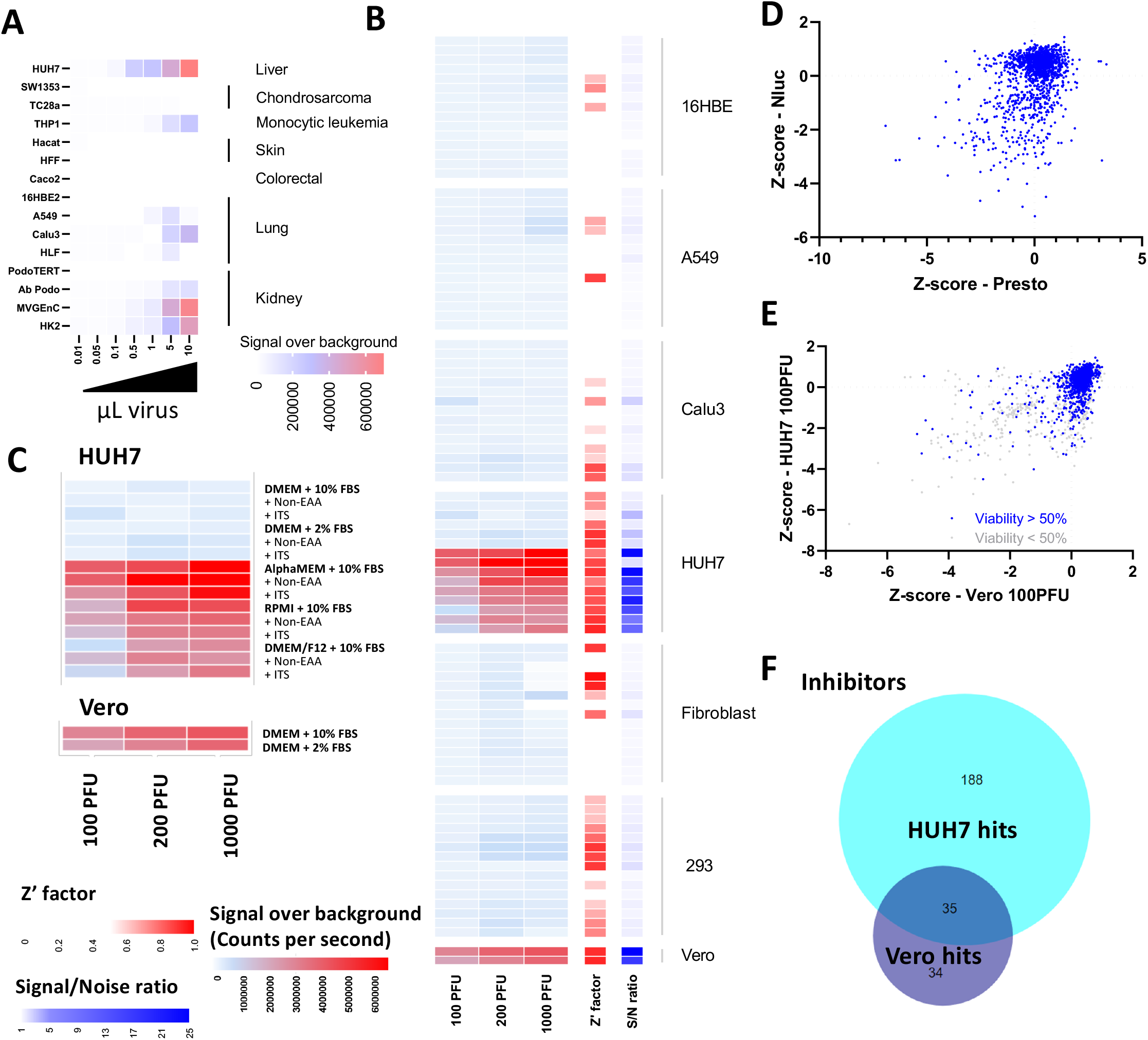
Replication of SARS-CoV-2-ΔOrf7a-Nluc in human cell lines. A) SARS-CoV-2 replication in human cell lines. Five thousand cells per well were seeded and infected as in Figure 3A. All cells were grown in the recommended growth medium, detailed in Table 1. The NLuc activity after 3 days was used to assess the degree of virus replication. NLuc activity above the baseline activity is shown as a heat map. Each box is the average of 2 biological repeats of each cell line at each virus dose, with the same findings observed when 2000, or 10,000 cells were seeded. B) Lung epithelial, fibroblasts, HUH7 and 293T cell lines were grown in 15 different growth medium compositions. NLuc activity over background three days after infection is shown. Each row represents a different growth medium as indicated in C). Each box represents 4 replicate measures of replication. Similar findings were observed in n=6 biological repeats. Z’ factors were calculated for each virus dose using ‘virus only’ wells as negative controls, the average Z’ factor for each growth medium is shown. The signal-to-noise ratio of infected cells relative to ‘virus only’ wells are also shown. C) NLuc activity above background three days after infection in HUH7 cells. Each box represents 4 replicate measures of replication. Similar findings were observed in n=6 biological repeats. D) The effects of 1971 FDA-approved compounds on the replication of SARS-CoV-2-Δ Orf7a-NLuc (100 PFU/well, MOI 0.02) in HUH7 cells grown in alpha-MEM supplemented with 10% FBS. Five thousand cells were treated with 10 µM of each compound for 24 hours prior to infection with 100 PFU of virus. Virus replication progressed for 72 hours as in Figure 3A. E) Comparison of compounds identified by screening for HUH7 and Vero cells Z-scores for luciferase activity are shown F) Comparison and overlap of inhibitors identified in the compound screening for HUH7 and Vero cells.

As each cell line has a specific culture medium for optimal growth, we set out to test if different media would support cell growth and virus replication in lung epithelial cells, as this is the primary site of Sars-CoV2 infection. We used 15 different medium conditions to grow lung epithelial cells prior to infection with SARS-CoV-2-ΔOrf7a-NLuc; for comparison, HUH7 cells were also included. Changes in growth conditions improved assay performance in lung epithelial and fibroblasts but signal-to-noise ratios remained low. In contrast, for HUH7 cells, these optimized conditions significantly improved the 96 well format assay as indicated by Z’ factor >0.5 and signal-to-noise ratios >20 (**Fig. 4B and C**). Using the 1971 FDA-compound library we identified 223 compounds that suppressed SARS-CoV- 2 replication by greater than 85% whilst maintaining cell viability. We refined these hits by overlapping with positive hits from the screen in Vero cells (**Fig. 4E, Supplemental Table 5**). We identified 35 inhibitory compounds and 2 compounds that increased NLuc activity, that were common to the two screens (**Fig. 4F**). The intended clinical use or targets of the 35 inhibitory compounds included anti-virals, antibiotics, modifiers of dopamine and estrogen receptor activity, calcium ion channel inhibitors and HMG-CoA reductase inhibitors (**Fig. 5A**). The hits also identified vitamin D3, which is available over the counter. Multiple vitamin D related compounds were present in the screen and these too suppressed SARS-CoV-2-Δ Orf7a-NLuc although these did not all meet our criteria for both Vero and human cell lines (**Fig. 5B**).

**Figure 5:**
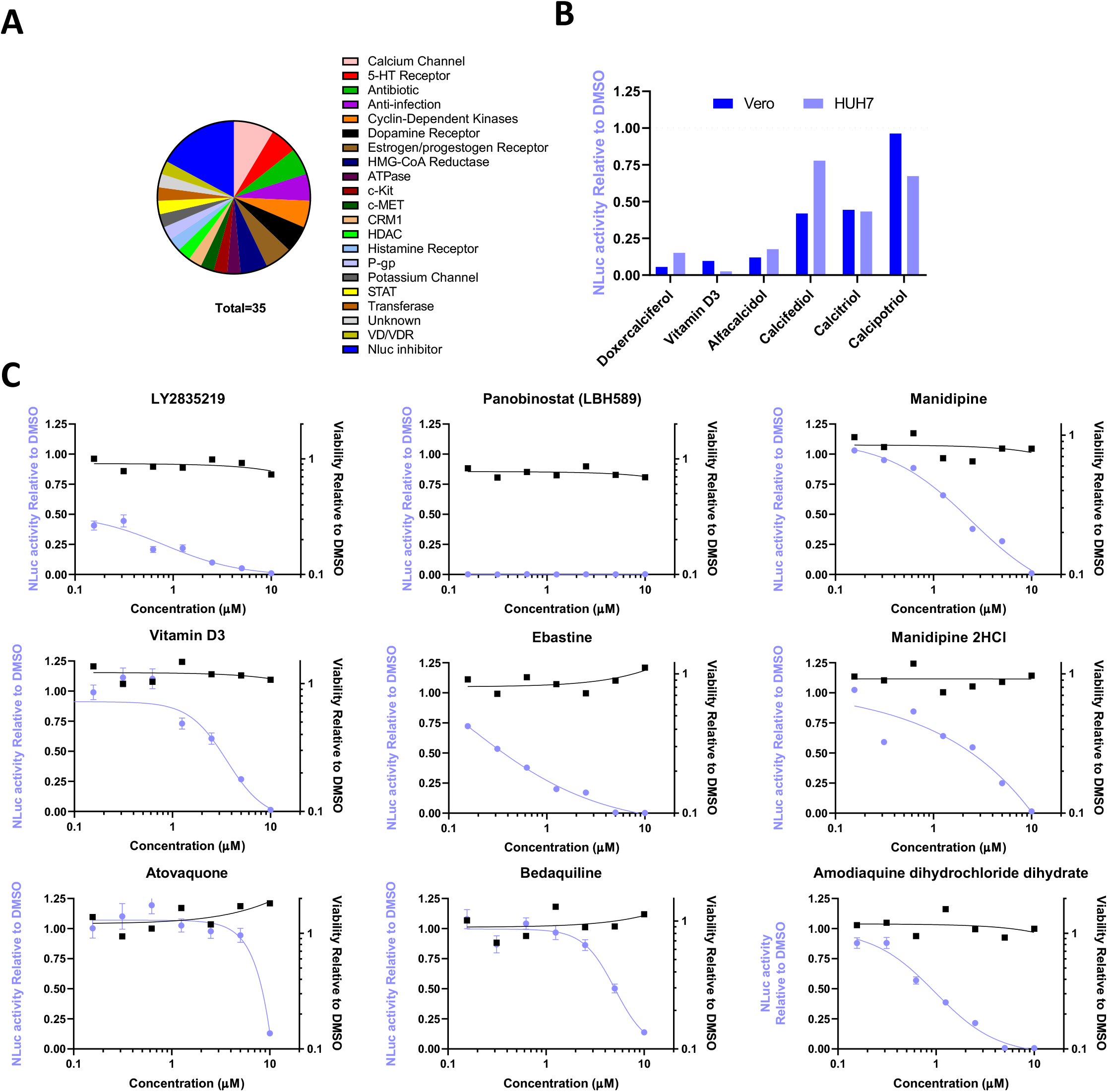
Dose response of SARS-CoV-2 inhibitors. A) Summary of the inhibitor class of the 35 compounds identified in both the Vero and HUH7 screens. B) Effects of vitamin D related compounds from the APExBIO DiscoveryProbe library on NLuc activity in the Vero and HUH7 screens. C) Dose responses to the 9 compounds in HUH7 cells. Cells were treated and infected as described in Figure 4D. NLuc activity relative to DMSO controls are shown in blue, and viability relative to DMSO are shown in black. N=2 independent repeats are shown.

**Table 1:** Approved uses of the 9 compounds of interest that inhibit SARS-CoV-2 infection and replication in HUH7 cells.

| Compound | HUH7 IC50 | Vero IC50 | Post-infection at IC50 | Approved Use | Target |
| --- | --- | --- | --- | --- | --- |
| Panobinostat | <0.2 μM | <0.2 μM | -25% | Multiple myeloma | HDAC inhibitor |
| LY2835219 | 0.2 μM | 1 μM | -40% | Abemaciclib for advanced breast cancer | CDK4/6 inhibitor |
| Manidipine | 2 μM | 7.5 μM | -50% | Anti-hypertensive | Calcium channel blocker |
| Manidipine 2HCl | 2.5 μM | 7.5 μM | -50% | Anti-hypertensive | Calcium channel blocker |
| Ebastine | 0.5 μM | 5 μM | -25% | Anti-histamine for allergic rhinitis and chronic idiopathic urticaria | H<sub>1</sub> receptor blocker |
| Atovaquone | 7.5 μM | 3 μM | -25% | antimicrobial (anti-malarial) | Selective inhibitor of parasite mitochondrial electron transport |
| Bedaquiline | 5 μM | 10 μM | -30% | pulmonary tuberculosis | mycobacterial ATP synthase |
| Vitamin D3 | 3 μM | 10 μM | -30% | Common supplement | Transcriptional/calcium regulation |
| Amodiaquine | 1 μM | 5 μM | -25% | antimicrobial (anti-malarial) | Inhibitor heme polymerase activity |

These initial hits were further tested to determine if treatment could also *prevent* replication in cells already infected with SARS-CoV-2-Δ Orf7a-NLuc. We infected Vero cells with SARS-CoV-2 for 24 hrs (which is sufficient time for the virus to begin replication) then added each of the 35 inhibitor compounds. The cells were incubated for 48 hrs (i.e. a total of 72 h.p.i) prior to bioluminescence being measured. The majority of the compounds had no impact on virus replication; but 9 of the 35 compounds reduced replication relative to DMSO controls (Table 1, **Supplemental Fig. 6**). The effective doses for these compounds were then determined in HUH7 cells to provide a future reference for selecting dose after comparison to pharmacokinetic data established in human trials (**Fig. 5, Table 1**). Whilst these are pre-clinical *in vitro* data, they demonstrate the efficacy of the compounds in reducing virus replication post infection and warrant further investigation to determine if these could ease the burden of the virus in patients.

## DISCUSSION

In this study we have shown that the SARS-CoV-2 virus infects and replicates in a range of human cells especially hepatocytes, kidney glomerulus, and proximal tubule cells of the kidney, and, that 9 drugs that have previously been shown to be safe in humans and approved by the FDA for clinical use are effective in inhibiting SARS-CoV-2 infection and replication.

The identification that liver and kidney cells are infection and replication competent for SARS-CoV-2 aligns with the observed liver and kidney abnormalities in patients with COVID-19. Liver comorbidities have been reported in 2-11% of patients with COVID-19, and 14-53% of cases reported abnormal levels of liver enzymes (Huang *et al*, 2020), with liver injury being more prevalent in severe cases (reviewed by (Zhang *et al*, 2020)). Another study reported that patients with chronic liver disease, especially African Americans, were at increased risk of COVID-19 (Wang *et al*, 2020). It has been suggested that liver damage in patients with COVID-19 might be caused by viral infection of liver cells, which is supported by the presence of SARS-CoV-2 RNA in stool (Yeo *et al*, 2020). On a similar note, acute kidney injury is a common complication of COVID-19 and has been associated with increased morbidity and mortality (reviewed by (Braun *et al*, 2020)). Our data showing SARS-CoV-2 infection and replication in kidney cells helps to explain kidney pathology associated with COVID-19.

RECOVERY (Group) has set out to identify therapeutics that could be re-purposed for treatment of COVID patients. For example, dexamethasone, which is a broad spectrum immunosuppressor, has been applied clinically to inhibit the destructive effects of the cytokine storm and shown to reduce mortality and decrease the length of hospitalization (Group *et al*, 2021), did not demonstrate any effect on viral replication in our study. It may be possible in the future to combine the therapeutics identified in our study with dexamethasone so that both the cytokine storm and virus replication are targeted.

Of interest, other therapeutics that have been tested in the RECOVERY trial include ritonavir and lopinavir (antiretroviral protease inhibitors), which failed to show clinical benefit in the treatment of COVID-19 (Group). These drugs were present in the DiscoveryProbe library used in our study but showed no significant effect on SARS-CoV-2 replication, in our assays.

The high costs and lengthy lead-in times associated with new drug development, make repurposing of existing drugs for the treatment of common and rare diseases an increasingly attractive idea (for review see (Pushpakom *et al*, 2019)). The approaches used include hypothesis driven, preclinical trials including computational (e.g., computational molecular docking) and experimental (e.g., biochemical or cell-based assays of drug interactions) methods, and evaluation of efficacy in phase II clinical trials. As noted by Pushpakom *et al*., of these three steps, the identification of the right drug for an indication of interest (in our case SARS-CoV-2 and COVID-19) is critical.

The nine drugs that we identified here have been approved for use in the treatment of a variety of diseases. Panobinostat, for example, is a HDAC inhibitor that blocks DNA replication, and has been used to inhibit cell growth in the management of cancer. Panobinostat had the strongest effect on limiting SARS-CoV-2 replication whilst maintaining cell viability, and completely blocked replication of SARS-CoV-2 at all doses tested (**Figure 5**); however, if cells were infected prior to treatment a more modest effect on replication were observed. This difference may be related to the recent observations that panobinostat can suppress ACE2 expression (He & Garmire, 2020), which would explain its beneficial effect prior to entry of the virus into cells and not thereafter. Abemaciclib (LY2835219) is another cell cycle inhibitor, suppressor of DNA replication and anti-cancer drug that emerged from the screen however these are likely to be detrimental to recovery from SARS-CoV-2 infection.

Atovaquone is of particular interest because it has been identified in other studies of SARS-CoV-2 in the context of COVID-19. It is a hydroxynaphthoquinone approved by NICE for the treatment of mild to moderate pneumocystis pneumonia and as a prophylaxis against pneumocystis pneumonia. It also has been used in combination with Proguanil (Malarone) as an antimalarial. The mechanism of action of atovaquone has been widely studied; it is a competitive inhibitor of ubiquinol, and against *P. falciparum*, it acts by inhibiting the electron transport chain at the level of the cytochrome bc1 complex (Fry & Pudney, 1992). A more recent study showed that atovaquone inhibits Zika and Dengue virus infection by blocking envelop protein-mediated membrane fusion (Yamamoto *et al*, 2020). Furthermore, in silico molecular docking strategies suggested potential binding of atovaquone to the SARS-CoV-2 spike protein (Farag *et al*, 2020; Ramirez-Salinas *et al*, 2020). These two mechanisms of action help to explain how atovaquone slows both infection and replication of SARS-CoV-2 in cells.

Our observation that bedaquiline inhibits SARS-CoV-2 replication supports evidence from an in silico repurposing study in which bedaquiline was proposed to be a promising inhibitor of the main viral protease of SARS-CoV-2 (Ferraz *et al*, 2020). The main protease cleaves pp1a and pp1ab polypeptides, which encode nonstructural proteins to form the replication-transcription complex (Snijder *et al*, 2006), and help explain how inhibitors to the main protease are effective in inhibiting replication of SARS-CoV-2 in Vero 76 cells (Ma *et al*, 2020). Manidipine (a calcium ion channel blocker approved for the use in treating hypertension) is proposed to reduce the activity of the main protease of SARS-CoV-2 (Ghahremanpour *et al*, 2020) and limit the availability of calcium ions, which are required for insertion of the coronavirus fusion peptide into the host lipid bilayers during viral fusion (Lai *et al*, 2017).

Our screen also identified ebastine (a second generation H1 receptor antagonist that has been approved for the treatment of allergic rhinitis and chronic idiopathic urticaria) and vitamin D3, which is a health supplement available over the counter. Ebastine is a second generation H1-antihistamine with proven efficacy for treating seasonal allergic rhinitis. However, histamine antagonists exhibit effects in addition to blockage of the histamine receptor. For example, ebastine blocks the release of anti-IgE-induced prostaglandin D2 (Campbell *et al*, 1996). The vitamin D receptor (VDR, a member of the nuclear hormone receptor superfamily) is proposed to be essential for liver lipid metabolism because its deficiency in mice protects against hepatosteatosis (Bozic *et al*, 2016). Once bound to VDR, vitamin D plays a major role in hepatic pathophysiology, regulation of innate and adaptive immune responses, and might contribute to anti-proliferative, anti-inflammatory and anti-fibrotic activates (reviewed by (Triantos *et al*, 2021)). Thus far, whether vitamin D supplementation reduces the risk of SARS-CoV-2 infection or COVID-19 severity is unclear (The Lancet Diabetes, 2021). Whilst vitamin D3 met the stringent cut-offs of 85% virus reduction, vitamin D2 and other Vitamin D related therapeutics also reduced virus replication but did not me*et al*l criteria across both Vero and HUH7 cell lines.

In conclusion, our study has identified compounds that are safe in humans and show effectiveness in reducing SARS-CoV-2 infection and replication in human cells, especially hepatocytes. Their potency in stopping SARS-CoV-2 replicating in human cells in the face of the COVID pandemic, warrants further study.

## MATERIALS AND METHODS

### Cell culture

Cell lines maintained in growth medium are shown in Supplementary Table 1.

### Generation of functional SARS-CoV-2 virus

DNA encoding the genome of SARS-CoV-2 and SARS-CoV-2-ΔOrf7a-NLuc were purchased from Vectorbuilder Inc. (Chicago, US). Transfection of virus encoding the DNA failed to generate replicative virus when electroporated into 293T cells. We therefore produced RNA molecules which encoded the virus *in vitro*. Briefly, virus encoding DNA (1 µg) was transcribed using T7 mMessenger mMachine (Thermo) with a GTP:Cap ratio of 2:1 used in a 20 µL reaction. In addition, RNA encoding the SARS-CoV-2 nucleocapsid was also generated by PCR using primers P1 and P2 (Supplementary Table 2). It has been reported that this aids the recovery of replicative virus (Insert ref 3). Viral RNA genomes (10 µl) and 2.5 µL nucleocapsid RNA were electroporated into 293T cells within the BSL3 laboratory. Viral RNAs were electroporated (5,000,000 cells, 1100 V, 20 ms and 2 pulses) and grown in T75 cm^2^ flasks and 24 well plates. Cells grown in 24 well plates for 24-120 hrs were used to monitor changes in NLuc activity.

### Virus production, maintenance and assessment of titer

Culture medium was collected from 293T cells 6 days after electroporation. This virus (P0) was used to infect cells of interest. As virus replication was slow in 293T cells, virus stocks were maintained by passage in Vero cells grown in DMEM supplemented with 2% FBS. Medium (1 mL) was used to infect Vero cells in order to generate P1 virus. Replication was assessed by measuring NLuc activity over 10 days. For subsequent passage of the virus, Vero cells in T75 cm^2^ flasks were infected with 2 mL of medium containing virus, after 3 days the medium was collected and passed through 0.45 µm filters using Luer-loc syringes.

To titre the virus, 200,000 Vero cells were seeded in 6-well plates overnight in growth medium. After removing growth medium Vero cells were infected with 200 µL of serially diluted virus containing medium at 37 °C. After 1 hour, wells were overlaid with 2 mL of 0.3% low melt agarose in 2xDMEM containing 1% FBS and grown for 3 days. After 3 days infected cells were fixed with 10% PFA overnight and then stained with crystal violet. Plaques were identified by imaging plates on BioDoc-It gel documentation system (UVP, Upland, US).

To detect viral RNA in medium, 0.25 mL of virus containing medium was collected 3 days post infection, 0.75 mL TriPure LS reagent (Sigma-Aldrich St. Louis, US) was added, and RNA isolated according to the manufacturer’s recommendations, in the final step RNA was dissolved in 15 µL DNAse/RNAse free water. For detection of SARS- COV-2 nucleocapsid transcripts in lung epithelial cells, 200,000 cells were infected at the indicated MOI for 3 days, monolayers were lysed directly in 1 mL Trizol (Invitrogen, Paisley UK). For assessing expression of ACE2, TMPRSS2 and NLP1 expression in lung epithelial cells RNA was isolated from the cells using Trizol and RNA isolated according to the manufacturer’s recommendations. For cDNA generation and real-time PCR we used conditions previously described (Pickard *et al*, 2019) using primers detailed in Supplementary Table 2.

### Electron microscopy

After 3 days infection with P4 virus, Vero cells were scrapped and pelleted. Cell pellets were fixed using 2.5% glutaraldehyde and 4% paraformaldehyde in 0.1 M cacodylate buffer for 24 h, washed in ddH_2_O three times, 30 mins for each wash. Cell pellets were incubated in freshly made 2% (vol/vol) osmium tetroxide and 1.5% (wt/vol) potassium ferrocyanide in cacodylate buffer (100 mM, pH 7.2) for 1 hr at room temperature. Samples were washed in ddH_2_O five times each for 3 minutes. Specimens were transferred to freshly made 1% (wt/vol) tannic acid in 100 mM cacodylate buffer (pH 7.2) for 40 mins at RT and washed in ddH_2_O five times for 3 mins each at RT. The specimen was incubated with 1% (vol/vol) osmium tetroxide in ddH_2_O for 30 minutes at room temperature and washed in ddH_2_O three times for 5 min each at room temperature. The specimen was then incubated with 1% (wt/vol) uranyl acetate (aqueous) at 4 °C for 16 hrs (overnight) and then washed in ddH_2_O three times for 5 mins each time at room temperature.

Specimens were dehydrated in graded ethanol: 30, 50, 70, 90% (vol/vol) ethanol in ddH_2_O for 10 mins at each step. Then samples were washed four times for 10 mins each time in 100% ethanol at room temperature. Samples were transferred to propylene oxide for 10 min at room temperature.

The specimen was finally infiltrated in a graded series of Agar100Hard in propylene oxide at room temperature: first for 1 hour in 30% (vol/vol) Agar100Hard, 1 hr in 50% (vol/vol) Agar100Hard then overnight in 75% (vol/vol) Agar100Hard, and then 100% (vol/vol) Agar100Hard for 5 hrs. After transferring samples to freshly made 100% Agar100 Hard in labelled molds and allowed to cure at 60 °C for 72 hrs. Sections were imaged on an FEI Tecnai12 BioTwin.

### NLuc activity assay

Vero cells grown in 24-well plates (Corning, 3526) were assayed for NLuc activity by adding 1 µL of coelenterazine (final concentration 1.5 µM). For 96 well formats, cell lines were seeded in white walled microwell plates (Nunc™ MicroWell™ 96-Well, Nunclon Delta-Treated, Flat-Bottom Microplate, Thermo Fisher Scientific, Paisley, UK# 136101). To measure NLuc activity, 0.5 µL coelenterazine was added per well (final concentration 3 µM). Light production was measured using filter cubes #114 and #3 on the Synergy Neo2 Multi-Mode Reader (Biotek), readings for each well were integrated over 200 ms with 4 replicate measurements per well (Gain 135 and read height 6 mm). For viability measurements, 2.5 µL Prestoblue (Thermo Fisher Scientific, Paisley UK) was added per well incubated for 10 mins before reading fluorescence at Excitation: 555/20 nm, Emission: 596/20 nm (Xenon flash, Lamp energy low, Gain 100 and read height 4.5 mm, 10 measurements per data point).

### Drug screens

The DiscoveryProbe FDA-approved library of 1971 compounds (L1021, APExBIO Boston, US) was prepared as follows. After thawing the library for 4 hours at room temperature the library was arrayed into 96 well plates at 1 mM in DMSO and stored at -20 °C. Stocks (1 mM) were thawed at room temperature for 2 hrs before compounds were added to cells.

For all drug screens and validation, 5000 cells were seeded in 50 µL of growth medium for 24 hrs in white walled microwell plates. DMEM containing 2% FBS was used for Vero cells and alphaMEM containing 10% FBS was used for HUH7 cells. The following day 0.5 µL of each compound (final concentration 10 µM) was added per well and incubated for 24 hours. Eighty-eight compounds were tested per plate and each plate contained the following controls: two untreated wells, two DMSO treated wells, and one well treated with 10 µM puromycin to kill cells. SARS-CoV-2- ΔOrf7a-NLuc virus was added to all wells. In addition, two wells did not contain any cells but were infected with the SARS-CoV-2-ΔOrf7a-NLuc virus as a measure of background NLuc activity. A final well which contained cells but were uninfected were also included. Twenty-four hours after drug treatment, cells were infected with 100 PFU per well SARS-CoV-2-ΔOrf7a-NLuc virus in 50 µL of the indicated growth medium for 72 hrs. To assess virus replication and viability, 2.5 µL of Prestoblue was added to each well and plates incubated for 10 mins at 37 °C. Coelenterazine was then added to a final concentration of 3 µM. Plates were sealed prior to reading luciferase activity and viability as described above. Validation of hits were performed using the same procedures described for the drug screen.

### Study design and statistical analysis

To identify hit compounds, raw luciferase luminescence reads (*x*) were normalised relative to the virus-infected drug-untreated controls (*u*) and plate minimum read (*m*) on each plate by the formula:

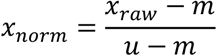

This meant that a normalized luciferase value of 1 implied no difference from untreated virus replication, and a value of 0 represented total inhibition of viral replication.

The PrestoBlue reads (*P*) were normalised relative to the virus-infected DMSO-treated controls (*d*) and plate minimum read (*n*) on each plate by the formula:

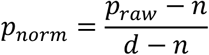

This meant that a normalized PrestoBlue value of 1 implied no difference in cell viability from DMSO-treated virus infected cells, and a value of 0 representing maximal reduction in cell viability.

Z-scores were calculated relative to the mean log_2_(fold change) for each plate.

Compounds were categorized as either inhibitors or enhancers of NLuc-SARS-CoV-2 activity. Inhibitors were compounds where normalized NLuc-SARS-CoV-2 levels (*x*_*norm*_) was reduced such that *x*_*norm*_ <0.15, and cell viability as measured by normalised PrestoBlue levels (*P*_norm_) wasn’t affected by more than 50%, such that *P*_*norm*_ > 0.5.

Where indicated one-tailed Student’s T-test were performed to evaluate significance in changes to NLuc activity.

## Contributors

AP co-conceived the project, performed the experiments, interpretated the data, drafted the manuscript. BCC performed experiments and interpreted the data. JC performed experiments. RG performed experiments. YL performed experiments. KEK co-conceived the project, raised the funding, drafted the manuscript. All authors approved the final manuscript.

## Acknowledgements

The research was funded by Wellcome (110126/Z/15/Z and 203128/Z/16/Z). The authors thank Dr. Jennifer Cavet for providing assistance with working at containing level 3 and use of the facilities. The project was approved by the COVID-19 Rapid Response Group, the R3G Research Operations Group and the R3G Executive Committee at the University of Manchester. We would also like to thank staff and students at the University of Manchester for the provision of cell lines for this study: Dr Stuart Cain, Dr Jonathan Humphries, Dr Shiu-wan Chan, Dr Chris Smith, Mrs Rachel Compton, Dr Rogerio Almeida, Dr Sara Gago, Dr Andrew Higham, and Dr Jeremy Herrera. Also thanks to Mrs Nikki-Maria Koudis, Mrs Maryline Fresquet, Dr Bernard Davenport and Dr Richard Naylor from Professor Rachel Lennon’s laboratory for help sourcing kidney cell lines and T7 mMessenger mMachine reagents. We would also like to thank Thermo Fisher Scientific for their help in providing microwell plates for this study, with special thanks to Charlotte Connor and Claire Marshall. We would like to thank Miss Anna Pickard for her illustration of the coronavirus.

## Figure legends

**Supplementary Figure 1:**
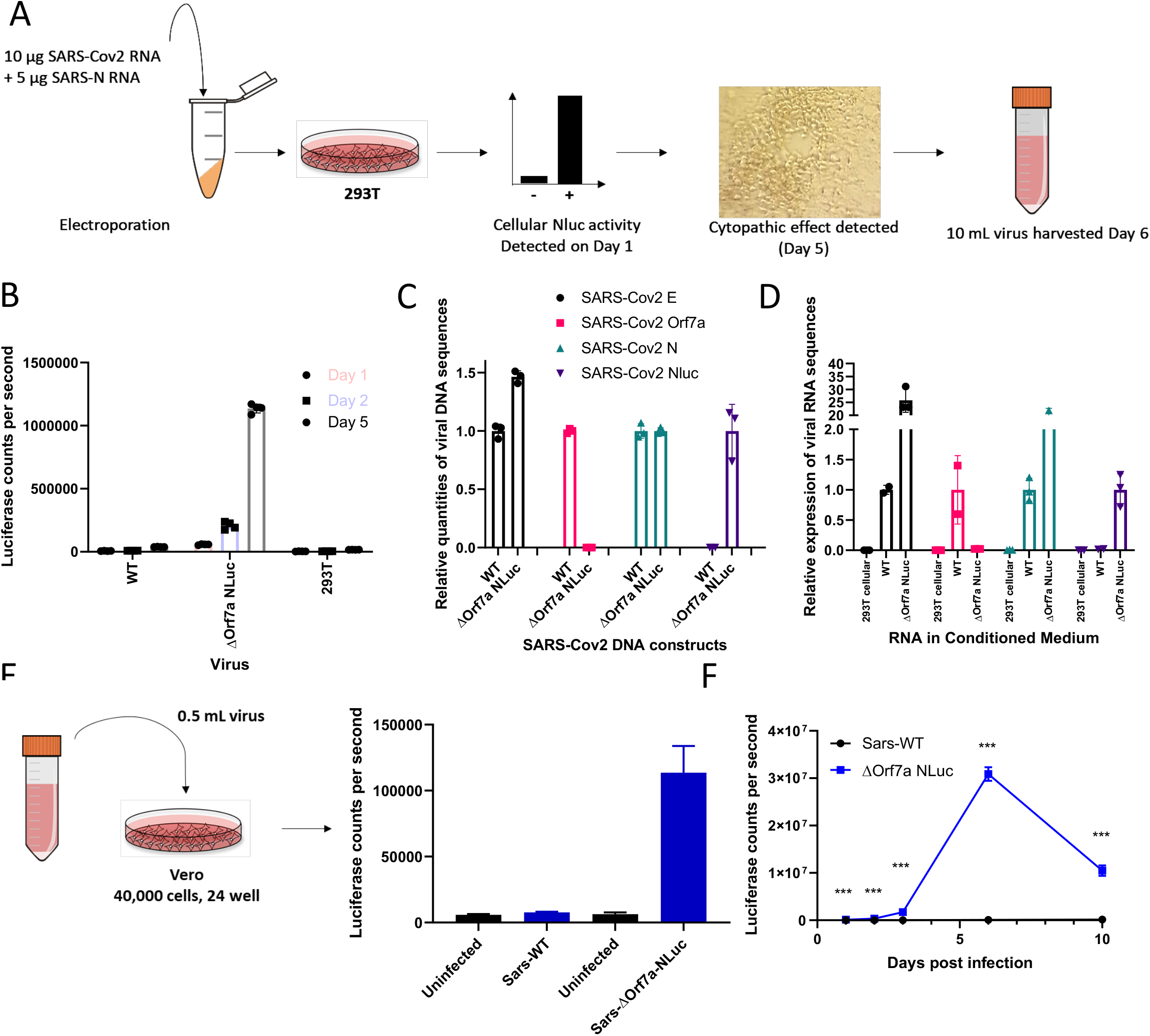
Recovery of replication competent SARS-CoV-2 from synthetic DNA constructs. A) Schematic for the recovery of SARS-CoV-2 virus particles from DNA encoding the wild type and NLuc modified SARS-CoV-2 genome. RNA was transcribed from the synthetic DNA constructs and electroporated into 293T cells, the cells were monitored daily for NLuc activity and evidence of the cytopathic effects of virus replication. Virus containing medium was collected 6 days post electroporation. B) NLuc activity in 293T cells electroporated with wild type and NLuc modified SARS-CoV-2 RNA transcripts. C) Real-time PCR primer validation using SARS-CoV-2 encoding DNA. NLuc coding sequences were inserted in place of the Orf7a gene. Primers designed to amplify sequences within the DNA constructs confirmed primer specificity. The number of cycles for each gene were normalized to those of the WT virus, or for NLuc, to Δ Orf7a NLuc. For each DNA construct the qPCR data was also normalized to the number of cycles obtained for the N gene (N=3 technical repeats). D) Real-time qPCR detection of viral RNAs in the medium of infected Vero cells 72 h.p.i. Vero cells were infected with Sars-CoV-2 WT (MOI 0.1) or Sars-CoV-2 ΔOrf7a-Nluc (MOI 1). E) Initial assessment of viral replication in Vero cells. Twenty four h.p.i. NLuc activity was readily detected in SARS-CoV- 2 ΔOrf7a-Nluc infected samples (N=3 replicate samples) F) NLuc activity in Vero cells infected with wild type SARS-CoV-2 or SARS-CoV-2 ΔOrf7a-Nluc modified virus. NLuc activity was significantly elevated in SARS-CoV-2 ΔOrf7a-Nluc infected cultures across all time points, *** represents p<0.001 in a Students T-Test, N=3 replicate samples. With prolonged culture NLuc activity within infected samples increased approximately 30-fold.

**Supplementary Figure 2:**
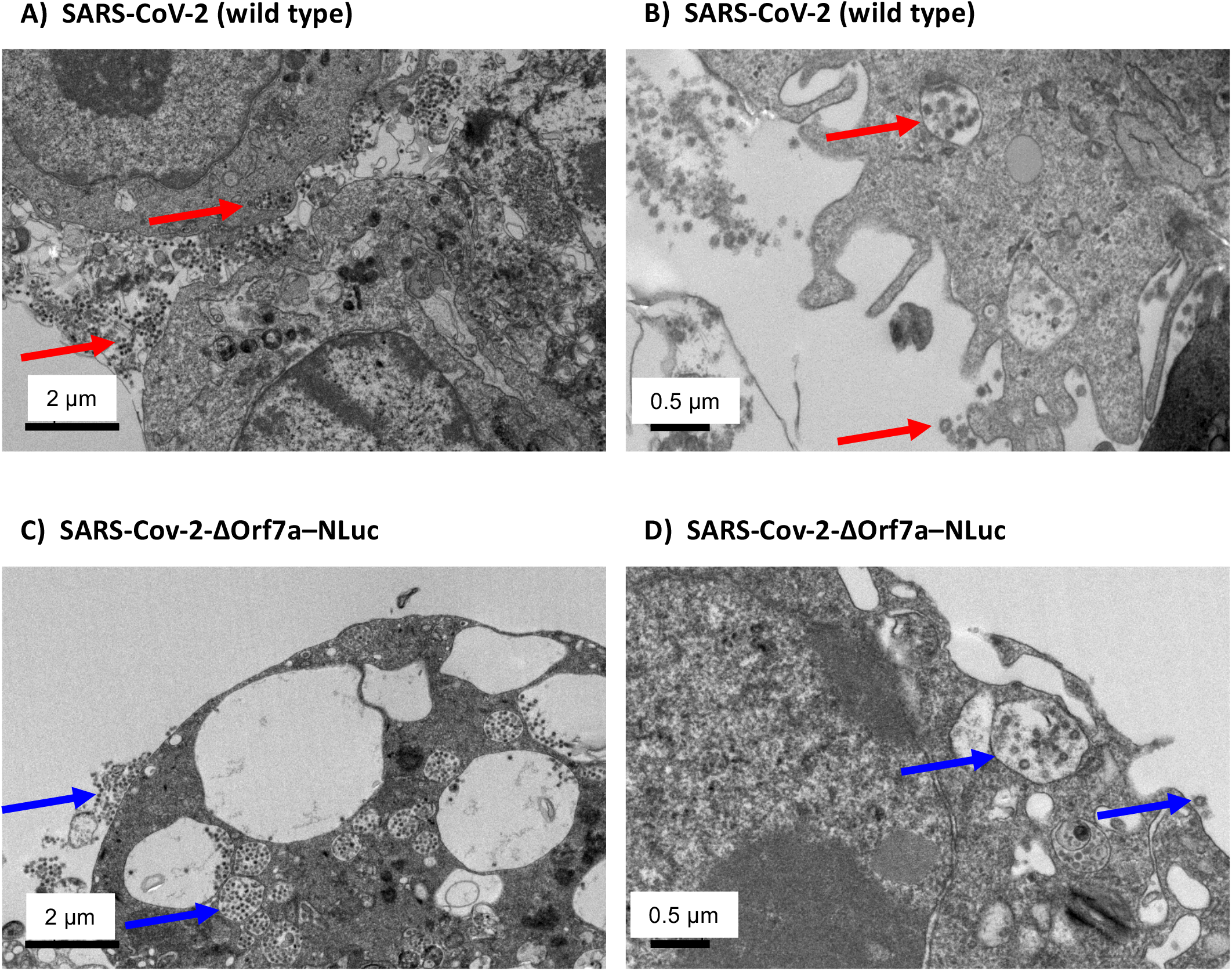
Detection of SARS-CoV-2 virus particles in Vero cells infected with passage 4 virus stocks. A) After 4 passages of recovered wild type SARS-CoV-2 virus particles in Vero cells, naïve Vero cells were infected and fixed 72 h.p.i. Viral particles could be observed inside and outside of Vero cells indicated by red arrows. Scale bar, 2 µm. B) Higher magnification image of an independent region of the sample in A. Scale bar, 0.5 µm. C) After 4 passages of recovered SARS-CoV-2-ΔOrf7a-NLuc virus particles in Vero cells, naïve Vero cells were infected and fixed 72 h.p.i. Viral particles could be observed inside and outside of Vero cells indicated by blue arrows. Scale bar, 2 µm. D) Higher magnification image of an independent region of the sample in C. Scale bar, 0.5 µm.

**Supplementary Figure 3:**
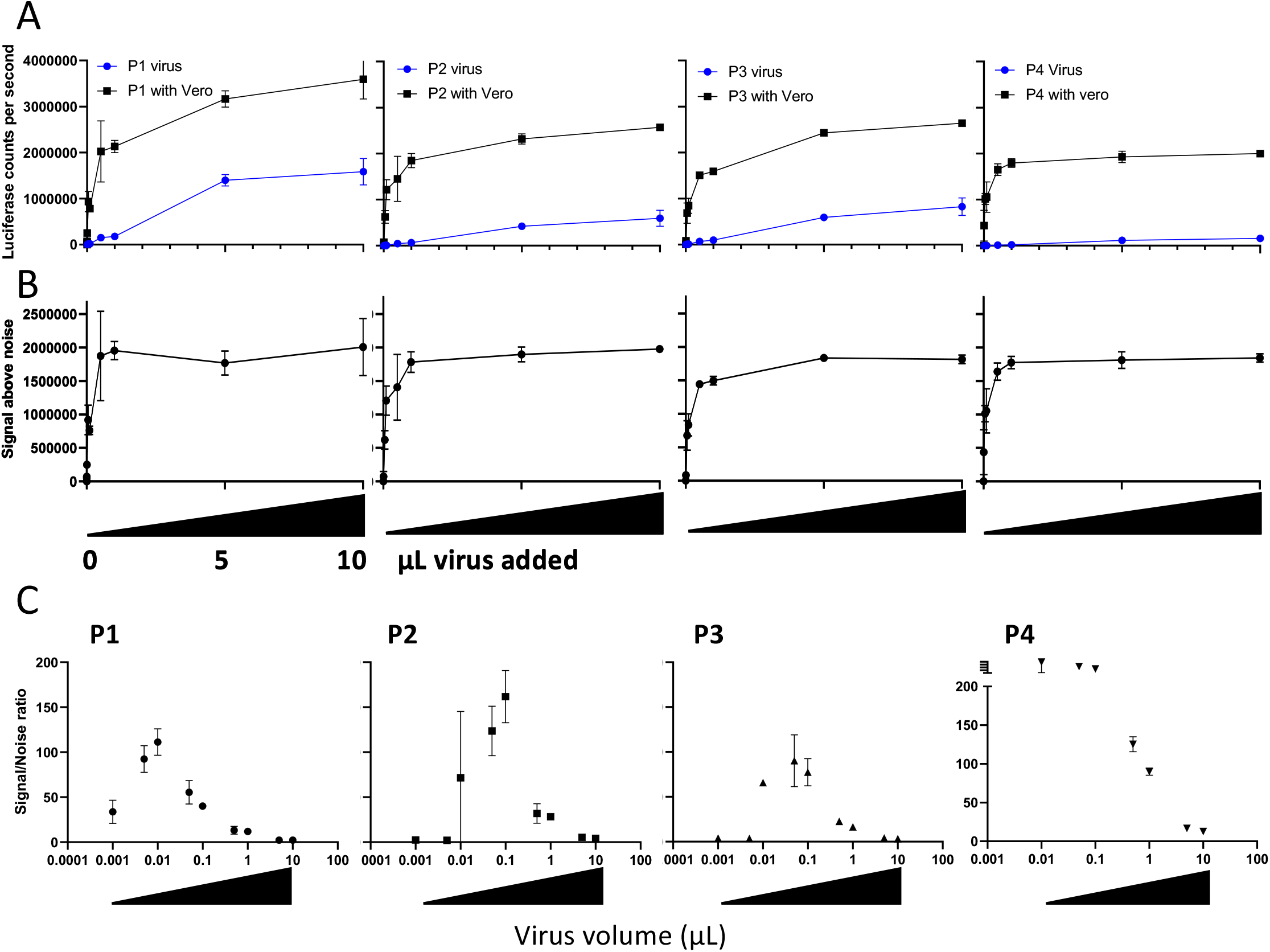
Replication assays for different passages of SARS-CoV-2-ΔOrf7a-NLuc virus. A) The plots show 5000 Vero cells infected with different volumes of virus harvested from repeat passages of the virus (P) in Vero cells. With increasing viral load, there is an increase in background as more NLuc activity is added to each well (Blue). The background signal arises when NLuc, expressed as the virus replicates, is released from infected cells that have lysed upon liberation of the virus particles. N=2 independent experiments. B) After subtracting the background noise observed in wells without cells (blue in A), the additional NLuc activity generated through viral replication showed comparable signals for each virus passage. N=2 independent experiments. C) The signal-to-noise ratio for each batch was used to identify the optimal conditions for virus replication for subsequent screening experiments.

**Supplementary Figure 4:**
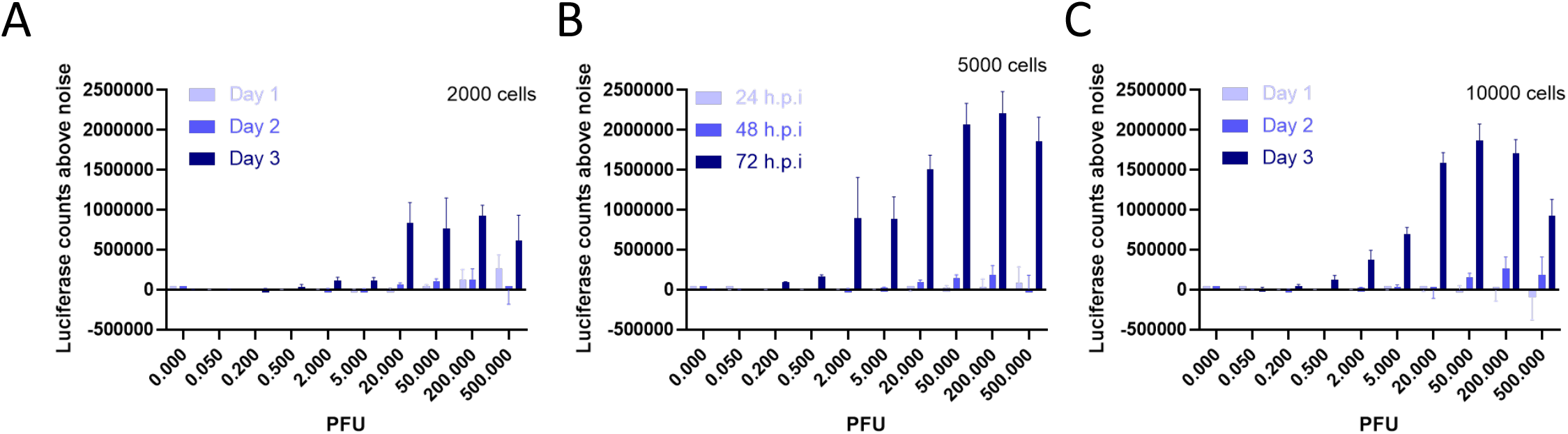
SARS-CoV-2-ΔOrf7a-Nluc replication optimisation. NLuc activity at different times after infection with increasing numbers of SARS-CoV-2 virus particles. Enhanced NLuc signals were observed 72 h.p.i. when either A) 2000, B) 5000 or C) 10,000 cells per well were used. Maximal signals were generated with 5000 cells per well, as seeding 10,000 cells tended to reduce NLuc activity. Five thousand cells were used for all subsequent assays.

**Supplemental Figure 5:**
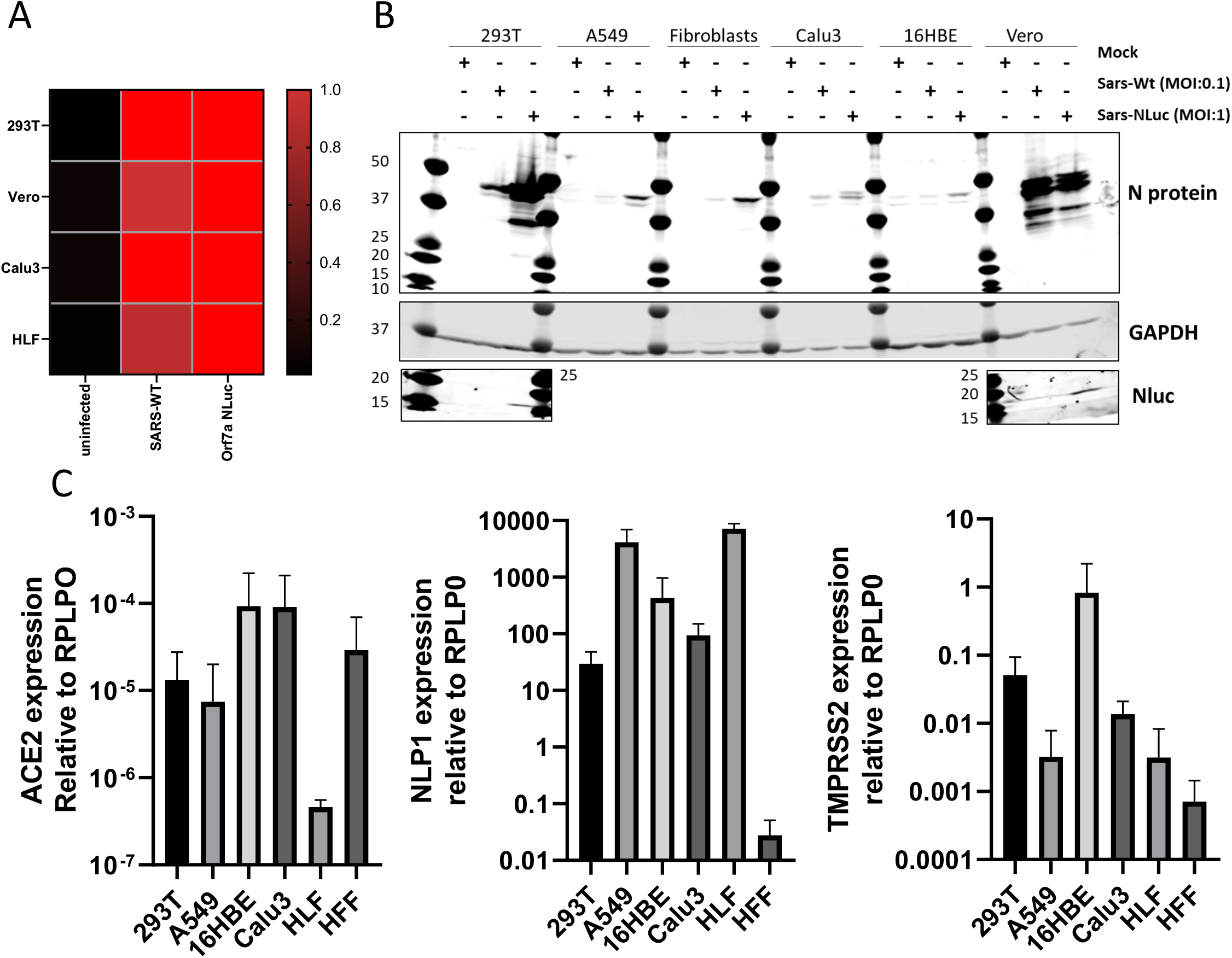
Detection of SARS-CoV-2 infection in lung epithelial cells. A) Real-time PCR detection of SARS-CoV-2 nucleocapsid (N) RNA in cells 72 h.p.i. Viral RNAs were readily detected in cells infected with WT or ΔOrf7a-Nluc virus particles. B) Western blot detection of SARS-CoV-2 nucleocapsid (N) protein in cells 72 hours post infection. Lower levels of the N protein are detected in fibroblasts and lung epithelial cells when compared to 293T and Vero cells suggesting the virus can infect but not replicate in these cell lines. C) Real-time detection of known mediators of SARS-CoV-2 entry into cells. ACE2, NLP1 and TMPRSS2 RNA levels are shown relative to RPLP0. Infectious virus in lung cell lines but no replication.

**Supplemental Figure 6:**
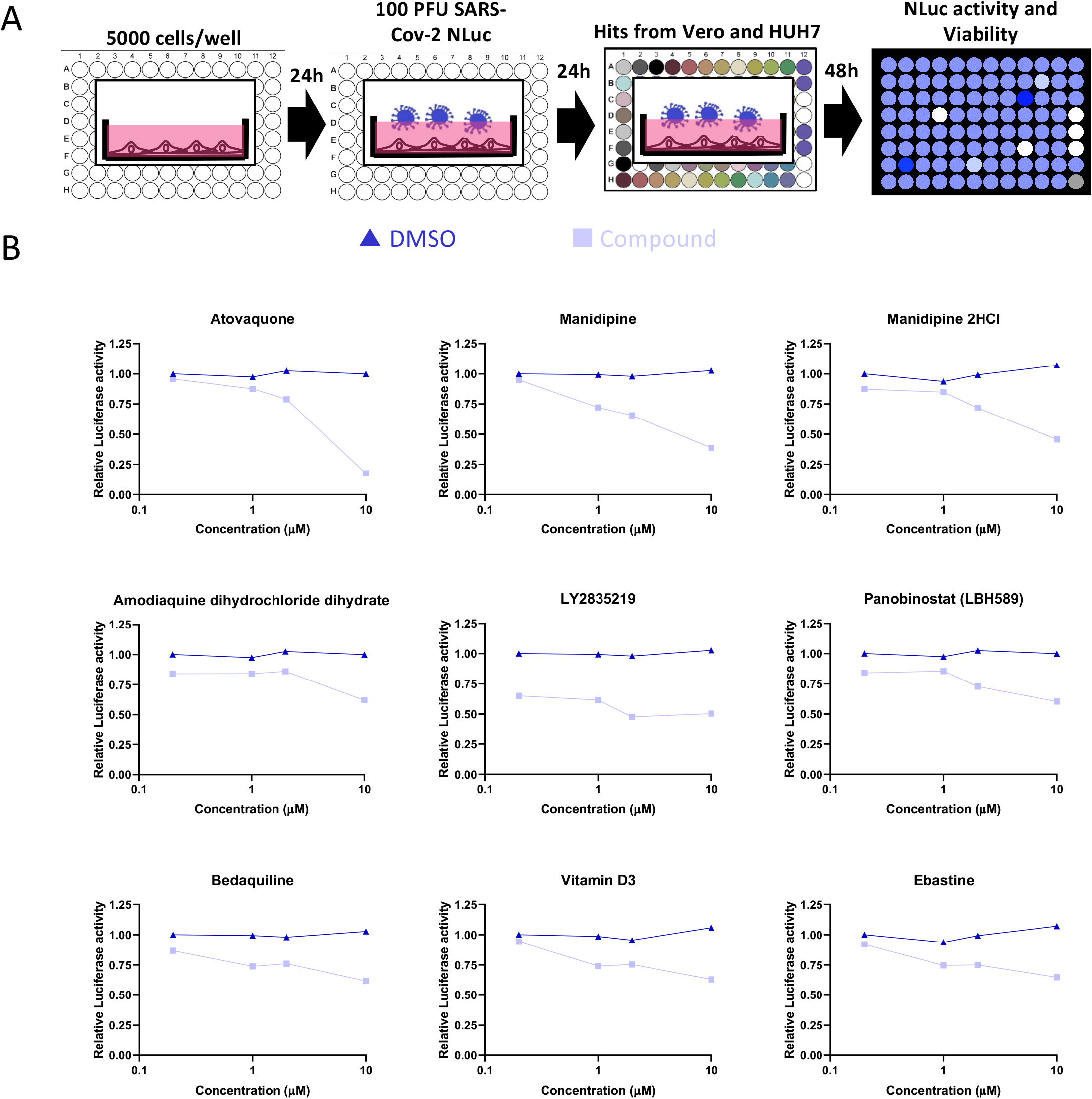
Effective compounds that suppress SARS-CoV-2 replication with treatment commencing after infection. A) Schematic for assessing whether any compounds that suppressed virus replication prior to infection could also affect replication post-infection. B) Fifty compounds identified to suppress SARS-CoV-2 replication in Figure 3 were assessed to identify dose-dependent effects on SARS-CoV-2 replication with treatment progressing after infection (as outlined in A). Nine compounds were found to suppress replication in a dose dependent manner.

**Supplementary Table 1:** Source and growth conditions for cell lines used in this study.

| Cell line | Growth medium | Source |
| --- | --- | --- |
| 293T | DMEM + 10% FBS | ATCC |
| HUH7 | DMEM + 10% FBS + NEAA | Accegen #ABC-TC0437 |
| THP1 | RPMI + 10% | Sigma-Aldrich, #88081201 |
| SW1353 | DMEM + 10% FBS | ATCC<sup>®</sup> HTB-94<sup>™</sup> |
| TC28a | DMEM + 10% FBS | Reference<sup>1</sup> |
| Hacat | DMEM + 10% FBS | Caltag Medsystems #T0020001 |
| HFF | DMEM + 10% FBS | Cascade Biologics/Thermo Fisher C0045C |
| Caco2 | DMEM + 10% FBS | ECACC #86010202 |
| 16HBE14o | AlphaMEM + 10% FBS | Reference<sup>2</sup> |
| Calu3 | AlphaMEM + 10% FBS | ATCC<sup>®</sup> HTB-55<sup>™</sup> |
| A549 | DMEM + 10% FBS | ATCC<sup>®</sup> CCL-185<sup>™</sup> |
| HLF (CCD-19Lu) | DMEM + 10% FBS | ATCC<sup>®</sup> CCL-210<sup>™</sup> |
| PodoTERT | PodoUP3 | Evercyte #PODO/TERT256 |
| MVGenC | Endothelial Basal medium-2 kit (without VEGF) | NeoBiotech #NB-11-0002 |
| Ab Podocytes | RPMI + 10% FBS + ITS | Reference<sup>3</sup> |
| HK-2 | DMEM:F12 + 10% FBS | ATCC<sup>®</sup> CRL 2190<sup>™</sup> |
| Vero | DMEM + 10% FBS | ATCC<sup>®</sup> CCL 81<sup>™</sup> |

FBS= Foetal bovine serum
NEAA=Non-essential amino acids
ITS= Insulin-Transferrin-Selenium
VEGF=Vascular endothelial growth factor.

**Supplementary Table 2:**
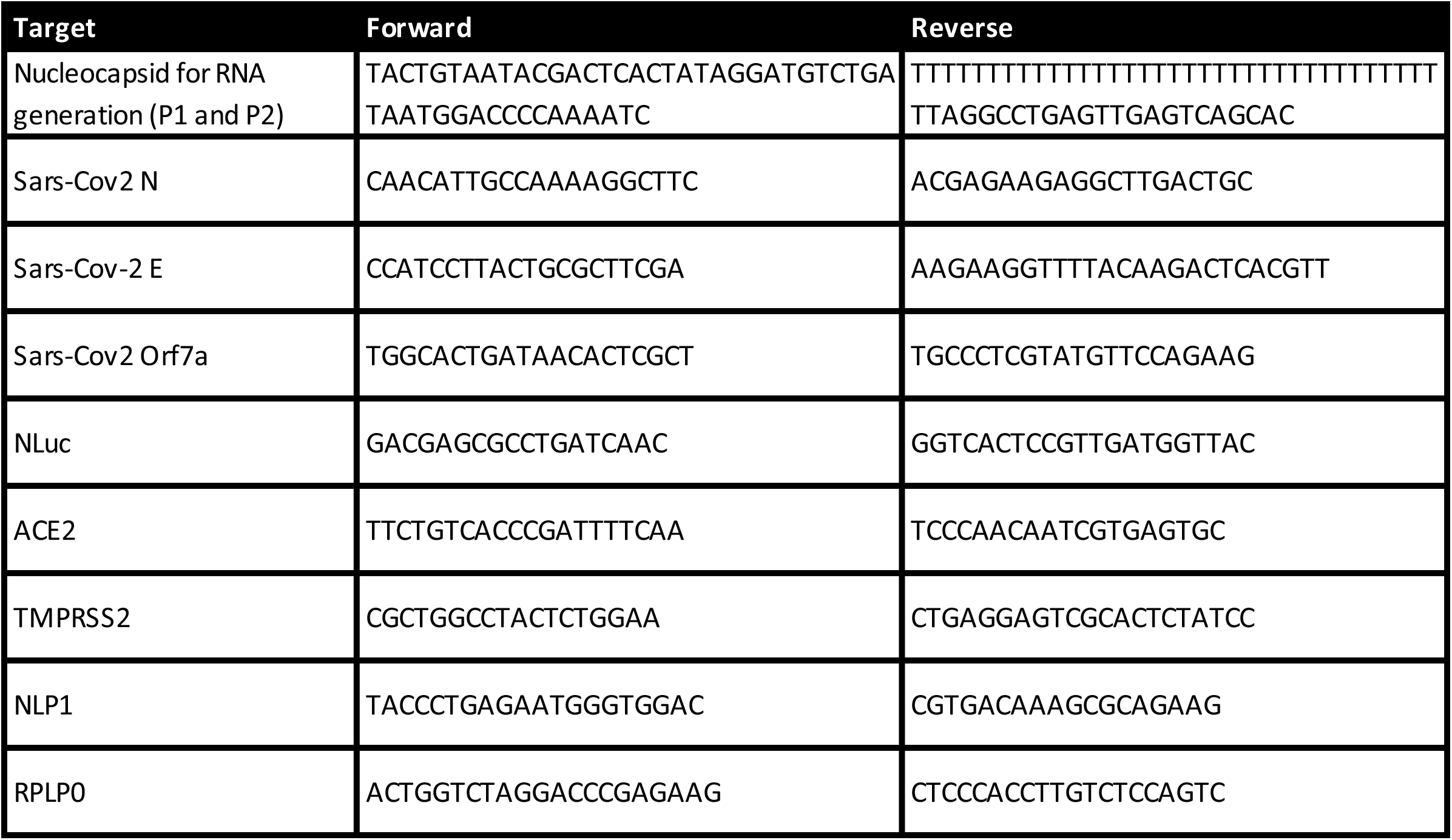
Primers used in this study

**Supplemental Table 3:**
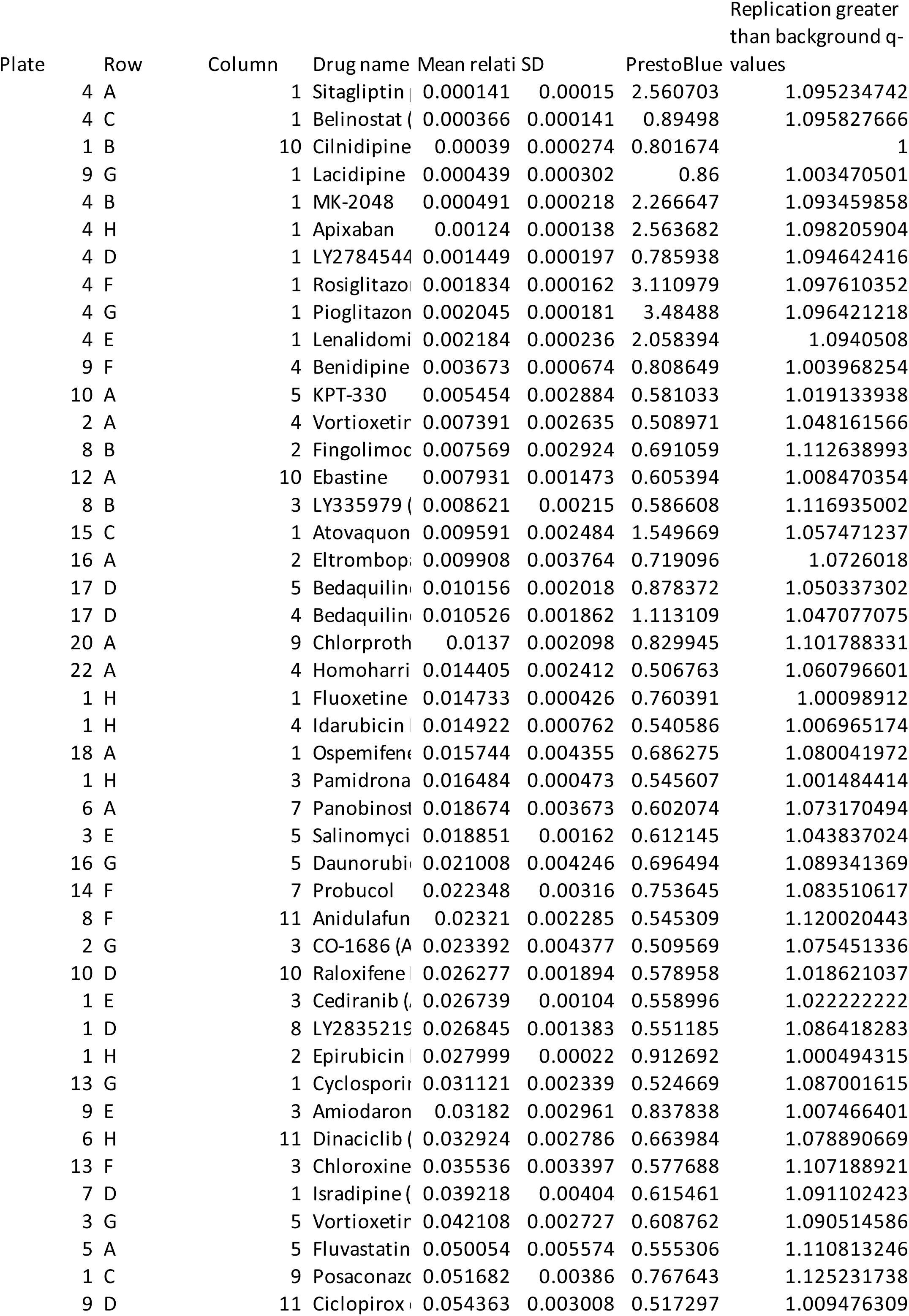
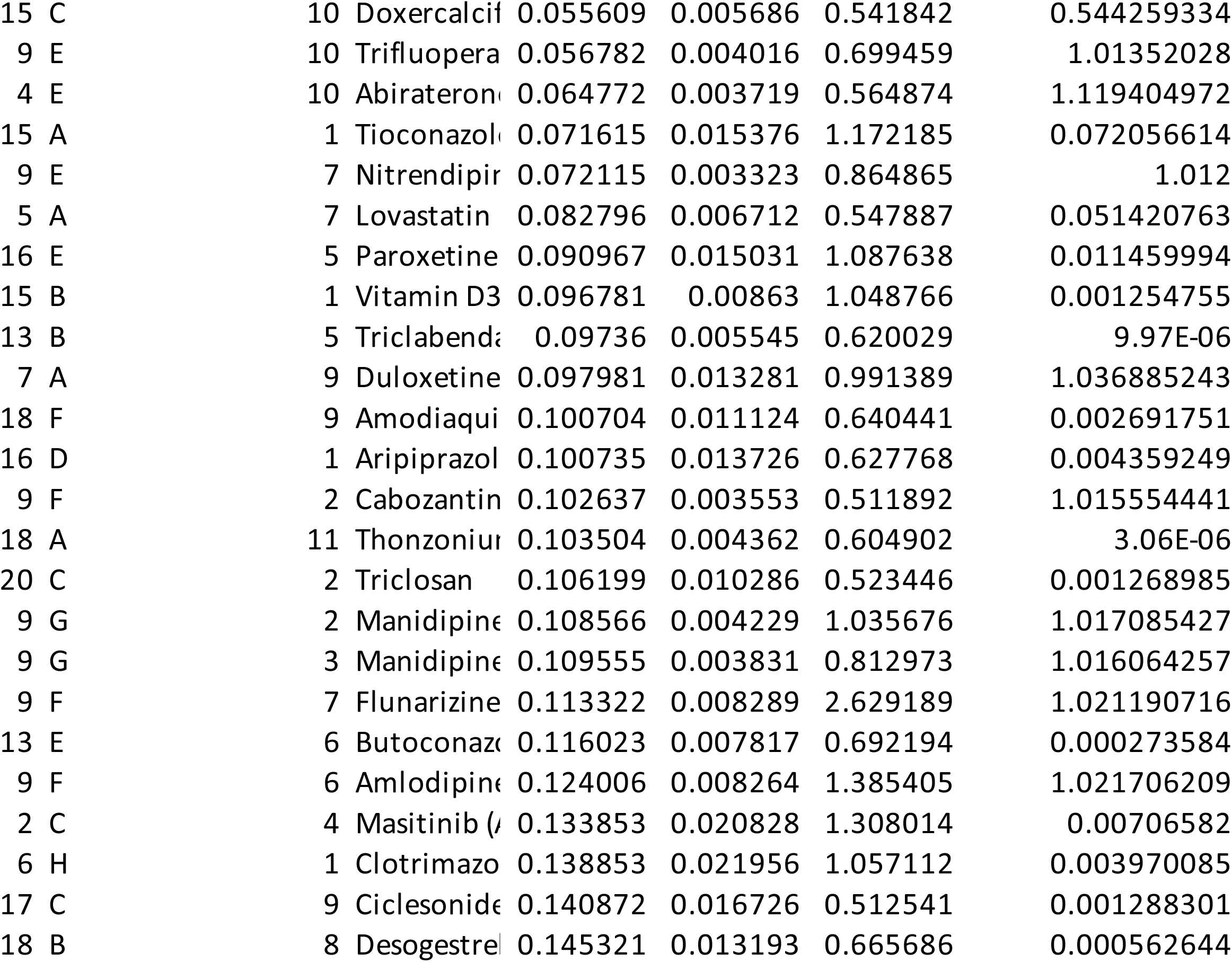
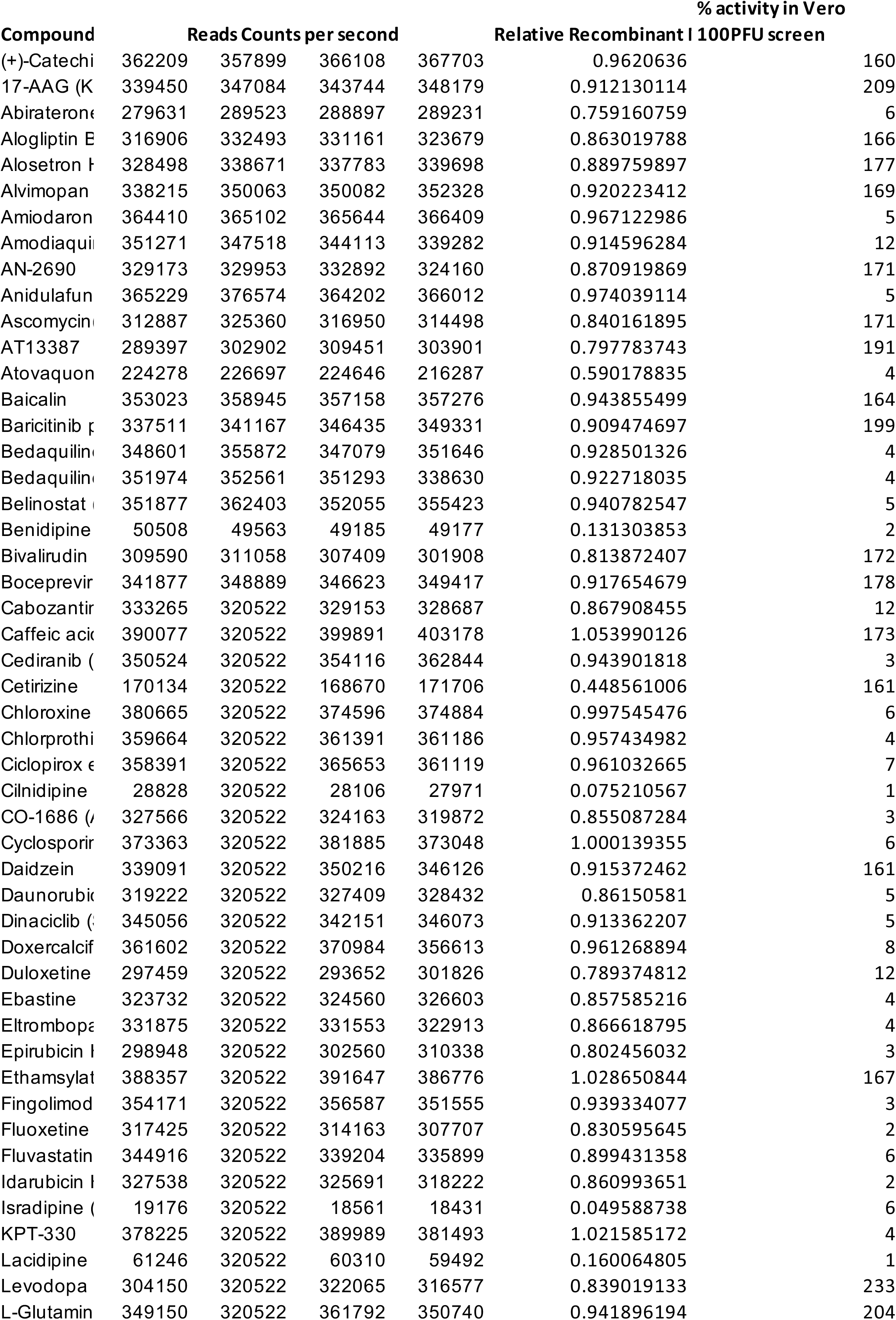
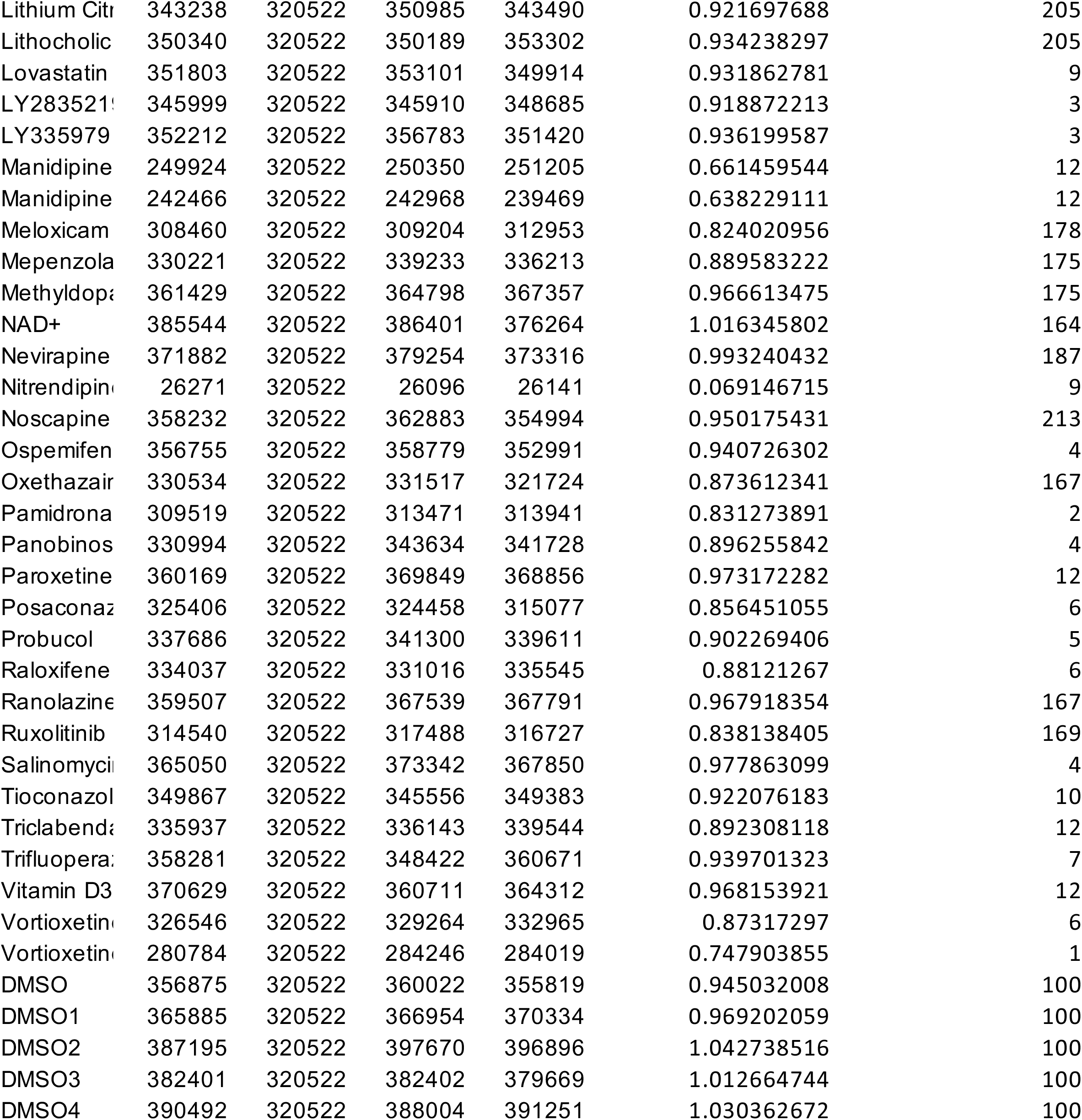
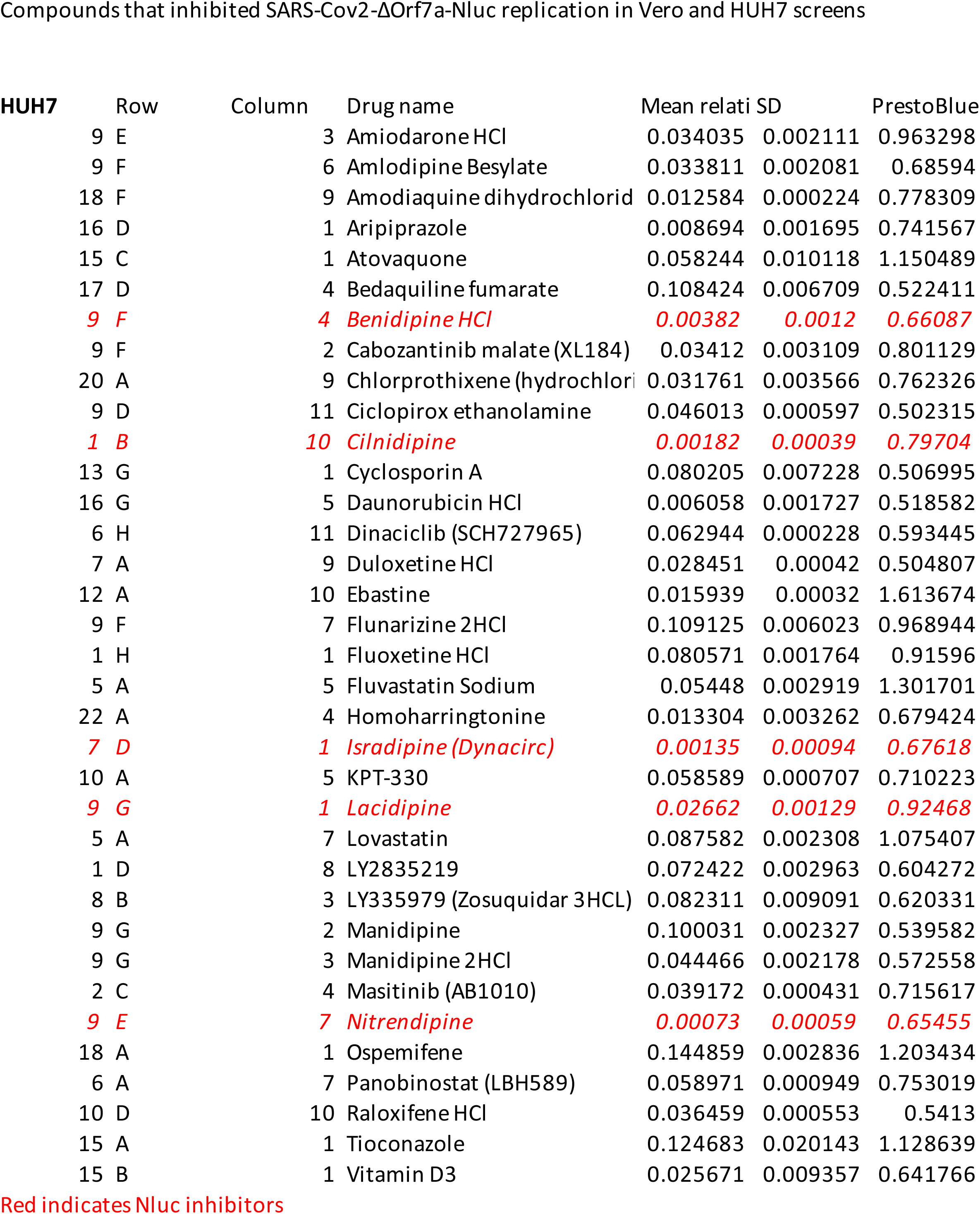
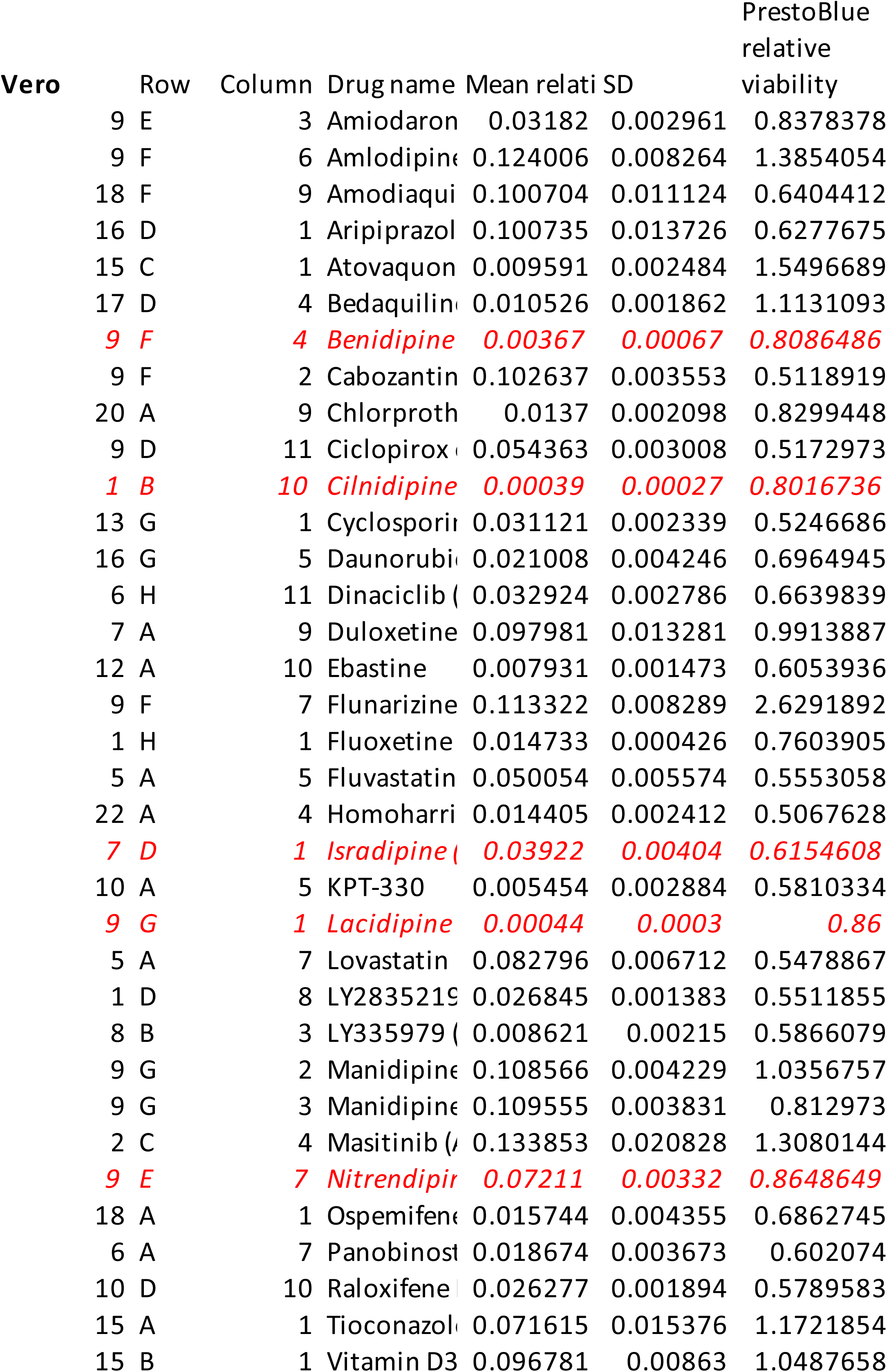
Inhibitors of Sars-Cov2-ΔOrf7a-Nluc replication in Vero cells

## References

Banerjee AK, Blanco MR, Bruce EA, Honson DD, Chen LM, Chow A, Bhat P, Ollikainen N, Quinodoz SA, Loney C et al (2020) SARS-CoV-2 Disrupts Splicing, Translation, and Protein Trafficking to Suppress Host Defenses. Cell 183: 1325–1339 e1321

Bozic M, Guzman C, Benet M, Sanchez-Campos S, Garcia-Monzon C, Gari E, Gatius S, Valdivielso JM, Jover R (2016) Hepatocyte vitamin D receptor regulates lipid metabolism and mediates experimental diet-induced steatosis. J Hepatol 65: 748–757

Braun F, Lutgehetmann M, Pfefferle S, Wong MN, Carsten A, Lindenmeyer MT, Norz D, Heinrich F, Meissner K, Wichmann D et al (2020) SARS-CoV-2 renal tropism associates with acute kidney injury. Lancet 396: 597–598

Calverley BC, Kadler KE, Pickard A (2020) Dynamic High-Sensitivity Quantitation of Procollagen-I by Endogenous CRISPR-Cas9 NanoLuciferase Tagging. Cells 9

Campbell A, Michel FB, Bremard-Oury C, Crampette L, Bousquet J (1996) Overview of allergic mechanisms. Ebastine has more than an antihistamine effect. Drugs 52 Suppl 1: 15–19

Cantuti-Castelvetri L, Ojha R, Pedro LD, Djannatian M, Franz J, Kuivanen S, van der Meer F, Kallio K, Kaya T, Anastasina M et al (2020) Neuropilin-1 facilitates SARS-CoV-2 cell entry and infectivity. Science 370: 856–860

Chatenoud L, Ferran C, Bach JF (1991) The anti-CD3-induced syndrome: a consequence of massive in vivo cell activation. Curr Top Microbiol Immunol 174: 121–134

Chen Y, Tao H, Shen S, Miao Z, Li L, Jia Y, Zhang H, Bai X, Fu X (2020) A drug screening toolkit based on the -1 ribosomal frameshifting of SARS-CoV-2. Heliyon 6: e04793

Chu H, Chan JF, Yuen TT, Shuai H, Yuan S, Wang Y, Hu B, Yip CC, Tsang JO, Huang X et al (2020) Comparative tropism, replication kinetics, and cell damage profiling of SARS-CoV-2 and SARS-CoV with implications for clinical manifestations, transmissibility, and laboratory studies of COVID-19: an observational study. Lancet Microbe 1: e14–e23

Fajgenbaum DC, June CH (2020) Cytokine Storm. N Engl J Med 383: 2255–2273

Farag A, Wang P, Boys IN, L. Eitson J, Ohlson MB, Fan Wea (2020) Identification of Atovaquone, Ouabain and Mebendazole as FDA Approved Drugs Tar-geting SARS-CoV-2 (Version 4). ChemRxiv

Ferraz WR, Gomes RA, A. SN, Goulart Trossini GH (2020) Ligand and structure-based virtual screening applied to the SARS-CoV-2 main protease: an in silico repurposing study. Future Med Chem 12: 1815–1828

Fry M, Pudney M (1992) Site of action of the antimalarial hydroxynaphthoquinone, 2-[trans-4-(4’-chlorophenyl) cyclohexyl]-3-hydroxy-1,4-naphthoquinone (566C80). Biochem Pharmacol 43: 1545–1553

Ghahremanpour MM, Tirado-Rives J, Deshmukh M, Ippolito JA, Zhang CH, Cabeza de Vaca I, Liosi ME, Anderson KS, Jorgensen WL (2020) Identification of 14 Known Drugs as Inhibitors of the Main Protease of SARS-CoV-2. ACS Med Chem Lett 11: 2526–2533

Group RC Lopinavir–ritonavir in patients admitted to hospital with COVID-19 (RECOVERY): a randomised, controlled, open-label, platform trial. The Lancet

Group RC A randomised trial of treatments to prevent death in patients hospitalised with COVID-19 (coronavirus). ISRCTN registry

Group RC, Horby P, Lim WS, Emberson JR, Mafham M, Bell JL, Linsell L, Staplin N, Brightling C, Ustianowski A et al (2021) Dexamethasone in Hospitalized Patients with Covid-19. N Engl J Med 384: 693–704

He B, Garmire L (2020) Prediction of repurposed drugs for treating lung injury in COVID-19. F1000Res 9: 609

Huang C, Wang Y, Li X, Ren L, Zhao J, Hu Y, Zhang L, Fan G, Xu J, Gu X et al (2020) Clinical features of patients infected with 2019 novel coronavirus in Wuhan, China. Lancet 395: 497–506

Jeon S, Ko M, Lee J, Choi I, Byun SY, Park S, Shum D, Kim S (2020) Identification of Antiviral Drug Candidates against SARS-CoV-2 from FDA-Approved Drugs. Antimicrob Agents Chemother 64

Lai AL, Millet JK, Daniel S, Freed JH, Whittaker GR (2017) The SARS-CoV Fusion Peptide Forms an Extended Bipartite Fusion Platform that Perturbs Membrane Order in a Calcium-Dependent Manner. J Mol Biol 429: 3875–3892

Li W, Moore MJ, Vasilieva N, Sui J, Wong SK, Berne MA, Somasundaran M, Sullivan JL, Luzuriaga K, Greenough TC et al (2003) Angiotensin-converting enzyme 2 is a functional receptor for the SARS coronavirus. Nature 426: 450–454

Ma C, Sacco MD, Hurst B, Townsend JA, Hu Y, Szeto T, Zhang X, Tarbet B, Marty MT, Chen Y et al (2020) Boceprevir, GC-376, and calpain inhibitors II, XII inhibit SARS-CoV-2 viral replication by targeting the viral main protease. Cell Res 30: 678–692

Maier HJ, Neuman BW, Bickerton E, Keep SM, Alrashedi H, Hall R, Britton P (2016) Extensive coronavirus-induced membrane rearrangements are not a determinant of pathogenicity. Sci Rep 6: 27126

Pickard A, Chang J, Alachkar N, Calverley B, Garva R, Arvan P, Meng QJ, Kadler KE (2019) Preservation of circadian rhythms by the protein folding chaperone, BiP. FASEB J 33: 7479–7489

Pushpakom S, Iorio F, Eyers PA, Escott KJ, Hopper S, Wells A, Doig A, Guilliams T, Latimer J, McNamee C et al (2019) Drug repurposing: progress, challenges and recommendations. Nat Rev Drug Discov 18: 41–58

Ramirez-Salinas GL, Martinez-Archundia M, Correa-Basurto J, Garcia-Machorro J (2020) Repositioning of Ligands That Target the Spike Glycoprotein as Potential Drugs for SARS-CoV-2 in an In Silico Study. Molecules 25

Riva L, Yuan S, Yin X, Martin-Sancho L, Matsunaga N, Pache L, Burgstaller-Muehlbacher S, De Jesus PD, Teriete P, Hull MV et al (2020) Discovery of SARS-CoV-2 antiviral drugs through large-scale compound repurposing. Nature 586: 113–119

Ruan Q, Yang K, Wang W, Jiang L, Song J (2020) Clinical predictors of mortality due to COVID-19 based on an analysis of data of 150 patients from Wuhan, China. Intensive Care Med 46: 846–848

Snijder EJ, van der Meer Y, Zevenhoven-Dobbe J, Onderwater JJ, van der Meulen J, Koerten HK, Mommaas AM (2006) Ultrastructure and origin of membrane vesicles associated with the severe acute respiratory syndrome coronavirus replication complex. J Virol 80: 5927–5940

The Lancet Diabetes E (2021) Vitamin D and COVID-19: why the controversy? Lancet Diabetes Endocrinol 9: 53

Thi Nhu Thao T, Labroussaa F, Ebert N, V’Kovski P, Stalder H, Portmann J, Kelly J, Steiner S, Holwerda M, Kratzel A et al (2020) Rapid reconstruction of SARS-CoV-2 using a synthetic genomics platform. Nature 582: 561–565

Touret F, Gilles M, Barral K, Nougairede A, van Helden J, Decroly E, de Lamballerie X, Coutard B (2020) In vitro screening of a FDA approved chemical library reveals potential inhibitors of SARS-CoV-2 replication. Sci Rep 10: 13093

Triantos C, Aggeletopoulou I, Thomopoulos K, Mouzaki A (2021) Vitamin D - liver disease association: Biological basis and mechanisms of action. Hepatology

Wang Q, Davis P, Xu R (2020) COVID-19 risk, disparities and outcomes in patients with chronic liverdisease in the United States. EClinicalMedicine

Wysocki J, Lores E, Ye M, Soler MJ, Batlle D (2020) Kidney and Lung ACE2 Expression after an ACE Inhibitor or an Ang II Receptor Blocker: Implications for COVID-19. J Am Soc Nephrol 31: 1941–1943

Xie X, Muruato A, Lokugamage KG, Narayanan K, Zhang X, Zou J, Liu J, Schindewolf C, Bopp NE, Aguilar PV et al (2020a) An Infectious cDNA Clone of SARS-CoV-2. Cell Host Microbe 27: 841–848 e843

Xie X, Muruato AE, Zhang X, Lokugamage KG, Fontes-Garfias CR, Zou J, Liu J, Ren P, Balakrishnan M, Cihlar T et al (2020b) A nanoluciferase SARS-CoV-2 for rapid neutralization testing and screening of anti-infective drugs for COVID-19. Nat Commun 11: 5214

Yamamoto M, Ichinohe T, Watanabe A, Kobayashi A, Zhang R, Song J, Kawaguchi Y, Matsuda Z, Inoue JI (2020) The Antimalarial Compound Atovaquone Inhibits Zika and Dengue Virus Infection by Blocking E Protein-Mediated Membrane Fusion. Viruses 12

Yeo C, Kaushal S, Yeo D (2020) Enteric involvement of coronaviruses: is faecal-oral transmission of SARS-CoV-2 possible? Lancet Gastroenterol Hepatol 5: 335–337

Zhang C, Shi L, Wang FS (2020) Liver injury in COVID-19: management and challenges. Lancet Gastroenterol Hepatol 5: 428–430

Zhang JH, Chung TD, Oldenburg KR (1999) A Simple Statistical Parameter for Use in Evaluation and Validation of High Throughput Screening Assays. J Biomol Screen 4: 67–73

Zhou P, Yang XL, Wang XG, Hu B, Zhang L, Zhang W, Si HR, Zhu Y, Li B, Huang CL et al (2020) A pneumonia outbreak associated with a new coronavirus of probable bat origin. Nature 579: 270–273

